# An fMRI meta-analysis of the role of the striatum in everyday-life vs laboratory-developed habits

**DOI:** 10.1101/2021.07.02.450904

**Authors:** Pasqualina Guida, Mario Michiels, Peter Redgrave, David Luque, Ignacio Obeso

## Abstract

The dorsolateral striatum plays a critical role in the acquisition and expression of stimulus-response habits that are learned in experimental laboratories. Here, we use meta-analytic procedures to contrast the neural circuits activated by laboratory-acquired habits with those activated by stimulus-response behaviours acquired in everyday-life. We confirmed that newly learned habits rely more on the anterior putamen with activation extending into caudate and nucleus accumbens. Motor and associative components of everyday-life habits were identified. We found that motor-dominant stimulus-response associations developed outside the laboratory primarily engaged posterior dorsal putamen, supplementary motor area (SMA) and cerebellum. Importantly, associative components were also represented in the posterior putamen. Thus, common neural representations for both naturalistic and laboratory-based habits were found in the left posterior and right anterior putamen. These findings suggest a partial common striatal substrate for habitual actions that are performed predominantly by stimulus-response associations represented in the posterior striatum. The overlapping neural substrates for laboratory and everyday-life habits supports the use of both methods for the analysis of habitual behaviour.

## Introduction

Repetition gradually enables human behaviour to become automated to the point where it can be executed without conscious thought and attention. Thus, over a lifetime, a vast array of sometimes sophisticated movements, thoughts, and motivations are assumed under automatic stimulus-response control. Typically, we refer to such behaviour as *habits*. Repeated associations between representations of specific stimuli and particular responses enable stimulus-evoked reactions to be enacted without conscious intervention (Haith and Krakauer, 2018). This allows us to perform established stimulus-response associations automatically, thereby allowing conscious attention to be directed to less predictable aspects of the world. An obvious example would be not having to think *how* to walk when consciously thinking about *where* to walk. Hence, the development of habitual routines enable us to perform predictable and less predictable tasks simultaneously – walking and talking (Sun, 2004).

A defining feature of habitual behaviour is once a stimulus-response association has been established, it proceeds independently of any change in the outcome value of the response (Dickinson et al., 1985; Miller et al., 2019). An example would be pressing the lift/elevator button taking you to the floor of your old office, (the previous stimulus-response outcome), rather than the new office. Here the lift is the stimulus while the habitual response would be pressing the old floor button. The move to the new office has devalued the outcome value of this response yet, when conscious attention is directed elsewhere, it is still performed. This so-called action slip example satisfies the outcome-devaluation criteria and would confirm its status as a habit. Automatic habitual control is often contrasted with conscious goal-directed processing where the action selected is determined by the value/appropriateness of the predicted outcome (Daw et al., 2005; Dickinson et al., 1985; Gläscher et al., 2010). Typically, before statistical regularities of a task are determined, adaptive goal-directed control is used flexibly to achieve the desired outcome at the expense of speed and efficiency. Consequently, early in instrumental learning, and in uncertain situations, flexible goal-directed control is specifically deployed for aspects of the task that cannot easily be predicted.

Human neuroimaging studies have revealed activation of the rostro-medial (associative) striatum during goal-directed instrumental learning, a pattern that gradually shifts to caudo-lateral (sensorimotor) regions when habitual control takes over (Brovelli et al., 2011a; de Wit et al., 2012a; Jankowski et al., 2009; Lehéricy et al., 2005a; Patterson and Knowlton, 2018; Tanaka et al., 2008; Tricomi et al., 2009). These findings concur with those described in non-human animals that demonstrate a similar involvement of the rostro-medial striatal territories early, and caudo-lateral regions later in the acquisition of instrumental tasks (Balleine and O’Doherty, 2010). Several neuropsychological studies including patients with striatal abnormalities provide converging evidence for a critical involvement of the caudal striatum in automatic habitual behaviour. The loss of dopaminergic input from the midbrain to the striatum in Parkinson’s disease is initially concentrated in the posterior putamen (Pineda-Pardo et al., 2022; Redgrave et al., 2010a). This selective loss seems to alter automatic moves such as arm swinging during walking or facial expressions in social interactions. More recently habitual components of typing (Bannard et al., 2019) and associative learning (Mi et al., 2021) seem to be reduced in Parkinson’s disease. In contrast, Huntington disease induces neural atrophy specifically in the head of the caudate (Vonsattel and DiFiglia, 1998) preventing the initial associations needed to integrate the repetition of stimulus-response associations into habits (Holl et al., 2012; Willingham and Koroshetz, 1993). Thus, evidence from experimental animal studies, functional imaging in healthy humans, and behavioural studies in human patients with selective dysfunction in different sub-regions of the basal ganglia, all point to the conclusion that conscious goal-directed and automatic habitual control are exercised by regionally segregated regions in the striatum.

A complication in the study of habits is the likely overlap with a wide range of other psychological concepts, including skills, automatic, routines, autonomous, inflexible, unconscious behaviour (West and Brown, 2013). We have already established that the formal definition of a habit is a stimulus-response association that proceeds automatically and is not modulated by outcome expectations (Dickinson et al., 1985). This perspective focuses on the stimulus evoked nature of habitual responses. Other definitions point more to the acquisition of habitual behaviours (Gardner, 2015), where the role of statistical regularities within tasks and the necessity of repetition are emphasised. Further, more recent analyses consider the way ultimate goals and motivation can influence habitual performance (Wood et al., 2021; Wood and Neal, 2007). However, due to the descriptive terminology used in different literatures, where the extent of conceptual overlap is unclear, inconsistencies and discrepancies between the previous studies of habitual behaviour should be expected (De Houwer, 2019). The purpose of the present meta-analysis was therefore, to determine the extent to which different forms of automatic stimulus-response behaviour engage the same or different neural substrates.

Until recently, most studies of habits have been in experimental settings where new stimulus-response relationships are learned in the laboratory. However, the relationship between these newly acquired stimulus-response associations and those acquired naturally over a lifetime when conducting everyday activities, is uncertain. It is equally important to understand how these important functional systems might malfunction, as reflected in brain disorders such as Parkinson’s disease (Bannard et al., 2019; Redgrave et al., 2010b), obsessive-compulsive disorder (Gillan, 2021), and drug addictions (Sjoerds et al., 2013), where dysfunctional patterns of habits are evident. Despite the obvious importance of habitual control, formal investigations of stimulus-response associations established in everyday-life are notably absent. An important step to remedy this would be to ask whether the neural circuits engaged by stimulus-response behaviour acquired in daily life are the same or different to those activated by newly learned habits in the laboratory. The specific purpose of the present investigation was therefore to compare the patterns of neural activation evoked by long-acquired habits brought into the laboratory with those newly acquired under formal experimental conditions.

Experimental research with animals has shown how instrumental learning occurs initially through goal-directed computations where behaviour is influenced by predicted outcome values and then later transitions into outcome-independent stimulus-response mappings (Balleine and O’Doherty, 2010). Specific tests have been devised to demonstrate the insensitivity of habitual responding to changes in outcome value or the stimulus-response relationship. For example, when instrumental performance is seen to persist after the outcome has been devalued, or the association between stimulus and response outcome degraded (see Box 1 for a summary), then the observed behaviour is considered under habitual rather than goal-directed control (Adams, 1982; Balleine and O’Doherty, 2010; Dickinson et al., 1985; Perez and Dickinson, 2020; for a review see Foerde, 2018). The formal procedures developed in animal studies have been imported into experiments investigating habits in humans (Luque et al., 2020; Tricomi et al., 2009; Valentin et al., 2007; Watson et al., 2014; Watson and de Wit, 2018).

### Box 1. Description of most used paradigms to assess everyday-life **(A)** behaviours and experimental learning **(B)** in fMRI contexts.

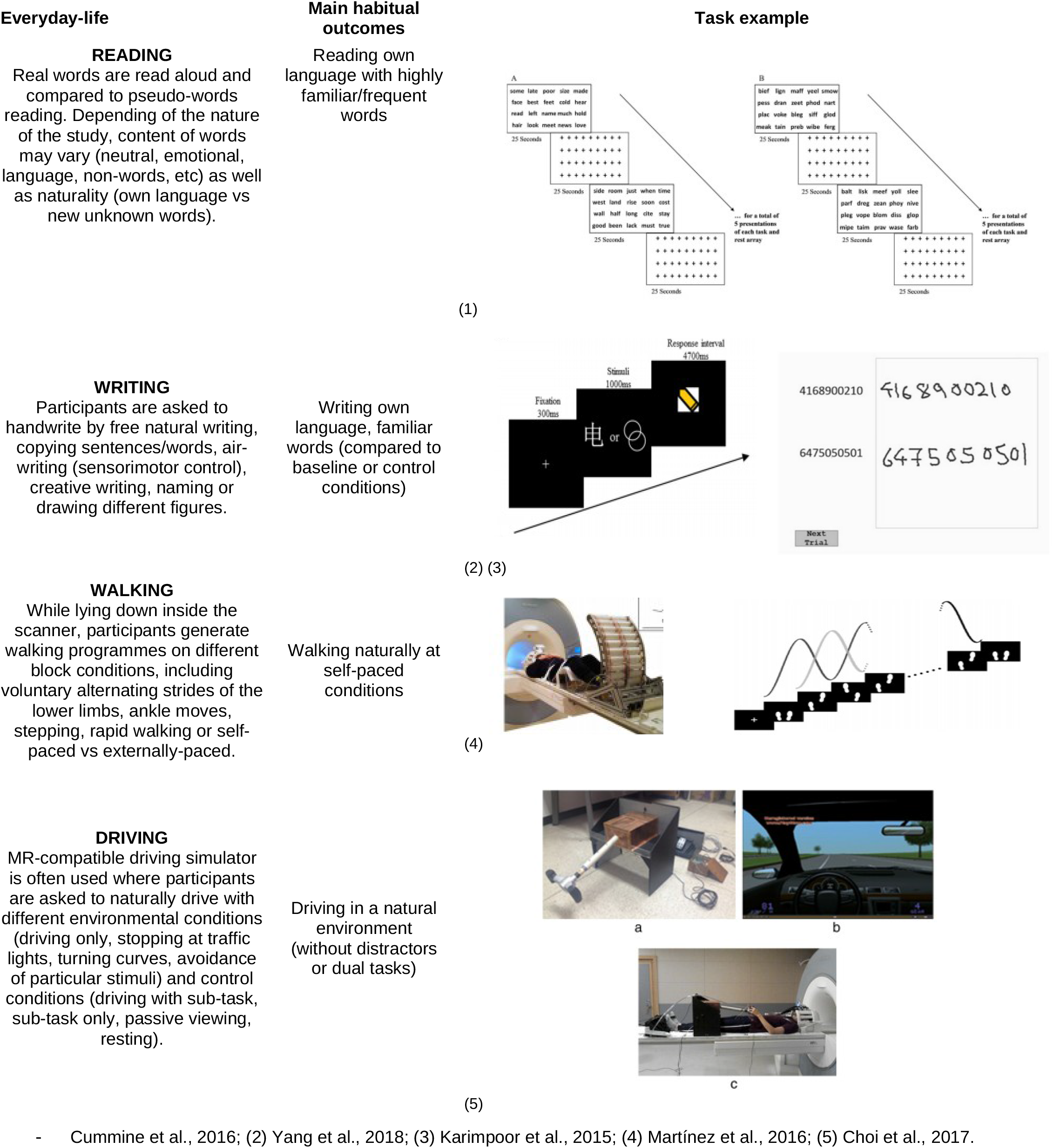
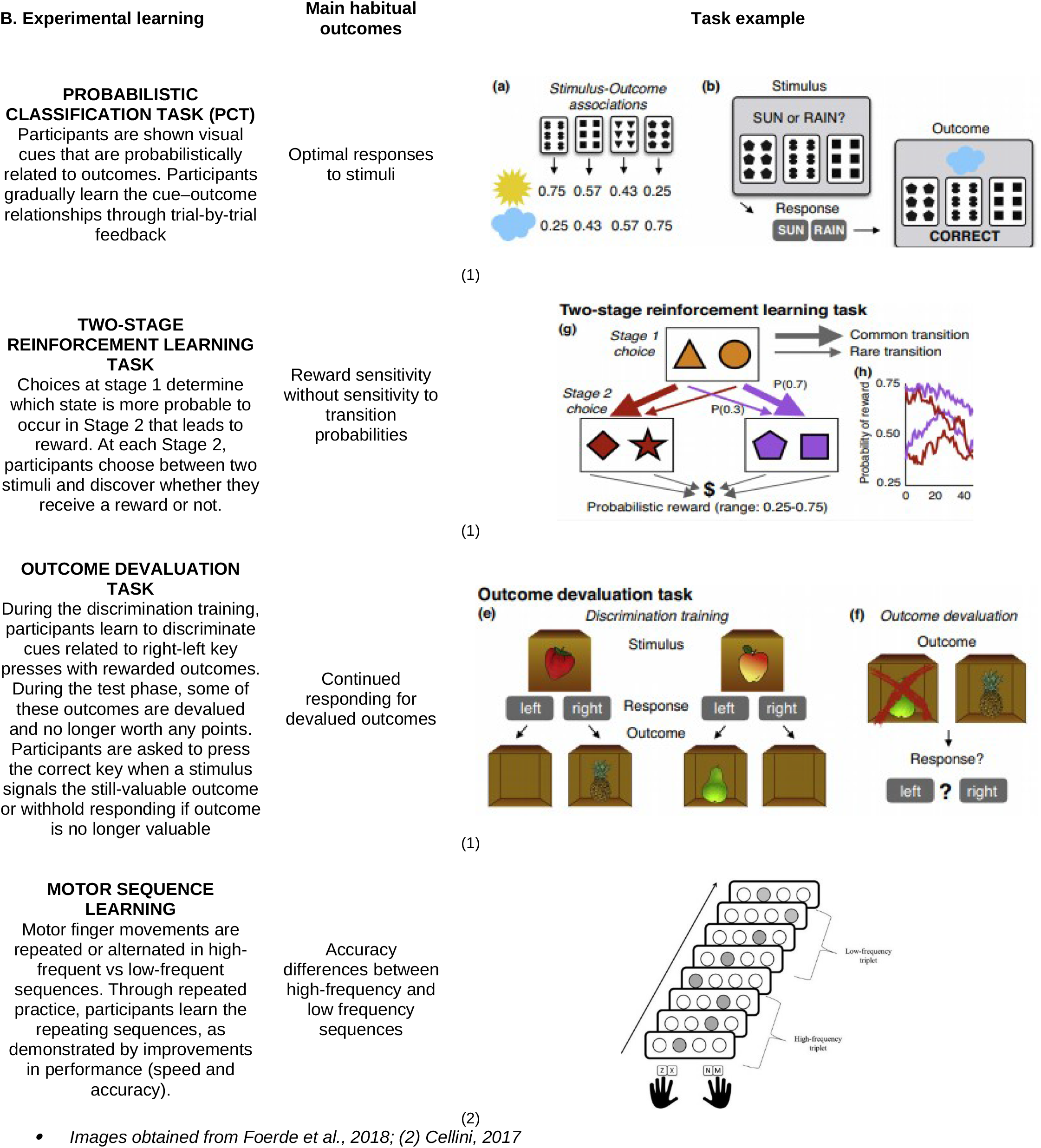

To date, most of the literature investigating the neural basis of human habits has relied on subjects developing new stimulus-response associations under formal <u>laboratory</u> conditions (Box 1B). This approach has provided important information about the transitions between goal-directed and habitual neural systems (Balleine and O’Doherty, 2010), and the factors (repetition, rewards or schedules of reinforcement) that promote the formation of habitual control (Dickinson et al., 1985; Omar D. Perez and Dickinson, 2020; Smith and Graybiel, 2016). However, this approach faces several challenges. Most pressing is the assumed equivalence of stimulus-response habits learned in the laboratory with those developed naturally in everyday-life. For example, one of the basic tenets of habitual behaviour is that the potency of stimulus-response associations should become stronger with repetition. It is therefore, not inspiring that several studies have failed to report a positive relationship between the amount of training and the strength of habitual responses measured in their experimental settings (de Wit et al., 2018; Pool et al., 2022.).

An alternative and increasingly important way forward to study habits in humans would be to have everyday habitual behaviour learned over a subject’s lifetime brought into the laboratory for investigation. Many behaviours in normal life including aspects of driving, eating, dancing, reading, talking or walking have significant stimulus-response components that can be performed automatically without thought while the person’s conscious attention is directed elsewhere (Wechsler et al., 2018; Worringer et al., 2019). Such components have been acquired through frequent repetition throughout a life-time of everyday trials. Such associations come to the laboratory fully formed and are independent of any new learning.

The principal challenge in studying everyday-life habits is getting subjects to express their long-established stimulus-response associations in laboratory settings. This is necessary so both the automatic behaviour and associated neural activity can be measured quantitatively. Consequently, investigators have chosen behaviours that have significant automatic stimulus-response components (Ceceli et al., 2020) and can be performed in the laboratory while both behaviour and fMRI BOLD signals are measured simultaneously. Examples of such tasks include reading, where comparisons are made between real words of different familiarity and emotional content, foreign words and pseudo-words (Cummine et al., 2016); writing and drawing (Lin et al., 2021; Yang et al., 2018) walking on a special apparatus (Martínez et al., 2018); and driving an MR-compatible driving simulator (Box 1A; Choi et al., 2017; Cummine et al., 2016; Huth et al., 2016; Karimpoor et al., 2015; Martínez et al., 2016, 2018; Oberhuber et al., 2013; Varotto et al., 2020; Yang et al., 2018). The specific question we addressed in the current study is whether the BOLD activity patterns reported for stimulus-response associations acquired in everyday life are the same, partially overlapping or separate from those described for recently established laboratory habits (Figure 1).

**Figure 1.**
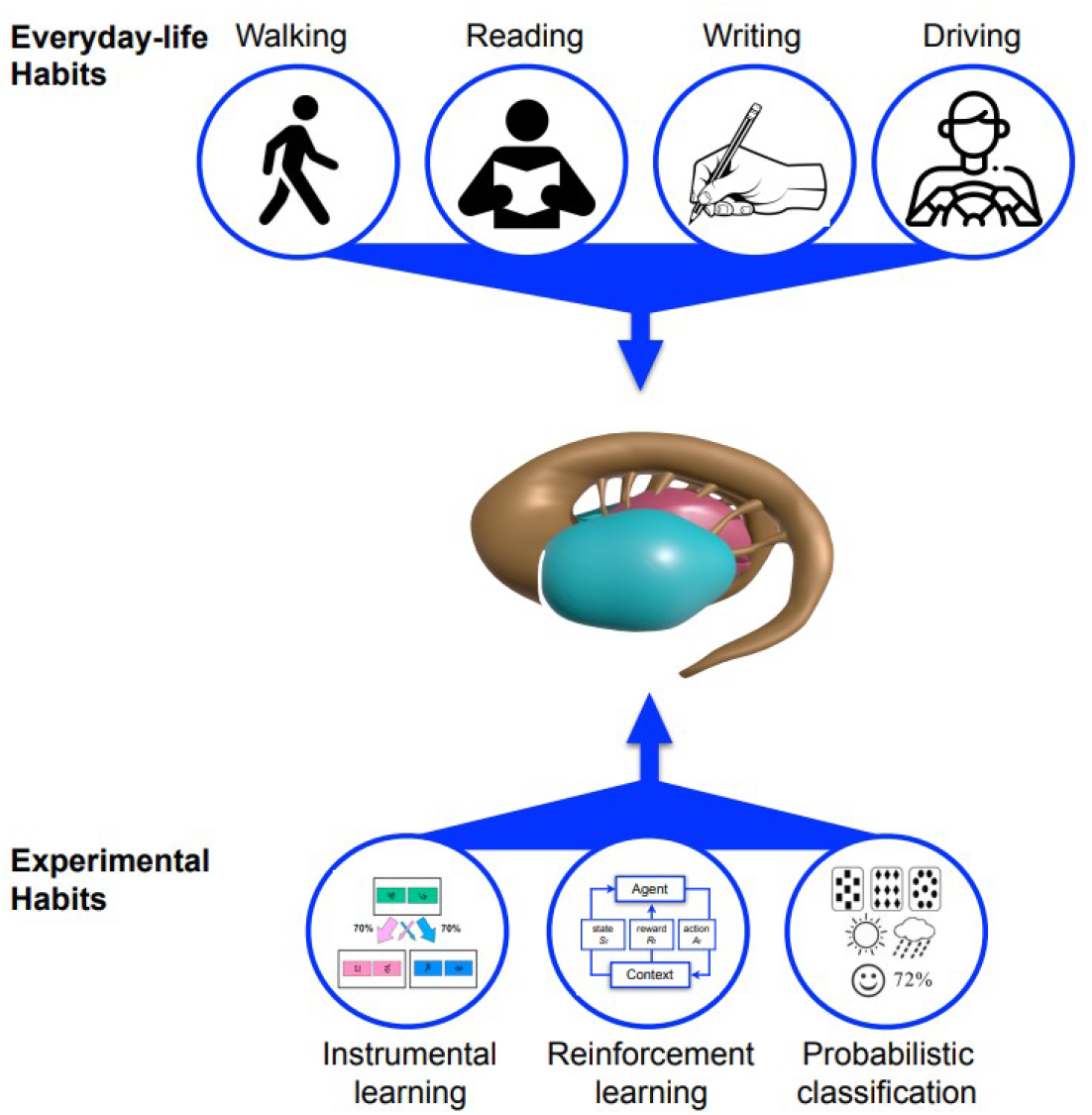
Hypothesis diagram on striatal role in both everyday-life and experimental habits. Activities part of daily life such as writing, reading, walking or driving were selected as everyday-life habits (see Box 1 for task measurements and details) to expect a critical striatal role in executing these habits forms. Similar striatal activities can be expected compared to experimental paradigms commonly used in the cognitive science literature.

To answer this question, we conducted a quantitative meta-analysis to investigate the neural substrates of everyday-life and experimental habits. This novel approach allowed a direct comparison of the respective neural signatures. Special interest was focused on the system-level circuits involving the basal ganglia (Figure 1). The resulting information provides important clues into the organisation of life-time habits against which pathologies of habit and the results of therapeutic interventions can be referenced.

## Methods

### Selection of studies

We conducted 2 independent searches in PubMed and Web of Science databases to identify fMRI studies investigating stimulus-response habits acquired: (1) in everyday-life; and (2) in experimental laboratories. The search of articles related to everyday-life habits was focused on natural behaviours expected to have significant automatic stimulus-response components: walking, driving, reading and writing. The selection of these activities was in part based on their potential link to fundamental basal ganglia functions (Figure 1) and were based on 3 additional criteria: (i) behaviours that are testable within an MR scanner; (ii) entail a strong component of stimulus-response learning; and (iii) acquired through lifetime repetition. A summary of typical measures used in these studies is presented in Box 1. In the search for laboratory based habits we included articles referenced by Patterson & Knowlton (2018) plus all new relevant articles published since their last search (June 22, 2017) and June 1, 2022. To enable direct comparison with, and the extension of the Patterson and Knowlton review (2018), we conducted literature searchs using the same key search terms and included the task categories – probabilistic learning, discriminative learning and sequence learning (see Box 1). We also included relevant MeSH terms available in Pubmed. This allowed us to find articles that do not include targeted keywords in the title or abstract but have proper MeSH terms linked to their metadata. The complete list of our search terms can be found in Supplementary Material. To set limits on inclusion, the articles were filtered further according to the following criteria:

(1) Neuroimaging modality and analyses: spatial coordinates from human brain fMRI reported in standardized stereotaxic space (MNI or Talairach space). Thus, other functional imaging methods (e.g., PET, EEG source imaging, etc) were excluded;

(2) Sample selection and signal sensitivity: healthy subjects over 18 years old were included, as were the healthy control participants within clinical studies with independent analyses. Since brain activity in sensorimotor-related regions of the basal ganglia can deteriorate with neurological conditions (Redgrave et al., 2010), we included studies only with healthy adult controls. Moreover, following other fMRI meta-analysis (Murphy and Garavan, 2004; Sescousse et al., 2013), we did not include studies with potential low sensitivity (less than 6 subjects);

(3) Whole-brain analysis: ROI analyses were excluded as they can be biased by the study hypothesis and may not report all significant regions;

(4) Computational models: for studies using computational model parameters (i.e., model-free and/or model-based parameters), we only selected those corresponding to the reward prediction error (RPE) and the 2-step probabilistic task (a task validated to assess the duality between habitual/goal-directed modelling (Daw et al., 2011). Model-based related parameters of any kind (e.g., state prediction error, SPE) were not considered in the analysis, as they are characteristics of goal-directed behavior;

(5) Type of studies and language: studies reporting original data and published in English peer-reviewed journals (other reviews and meta-analysis were excluded).

We then performed a second-step control using reference lists from already included articles. From the list of references, we first excluded articles whose title had no direct link to any of our inclusion criteria. From the remaining articles that met our inclusion criteria we found 17 that had investigated everyday-life habits and 14 articles that had investigated laboratory-developed habits. Since our second exclusion step involved manual, rather than online searching, we were able to include articles published before 2017 not included in Patterson and Knowlton (2018). These additional references were selected because: (1) we checked all references from the list of accepted papers, not only those in major review articles; and (2) our criteria allowed us to include studies that did not report putative habit-related activation of the dorsal striatum (caudate, putamen, or both). Figure S1 depicts the identification, screening, eligibility and selection stages we used to select studies for inclusion. Foci, scripts and statistical maps can be accessed in the Open Science Framework (https://osf.io/w5ftm).

### Contrasts of interest

The selected contrasts were based on the task condition reflecting the least difficulty level and less goal-directed task demands as well as comparisons linked to stimulus-response behaviours (see Table S1 and S2). Of interest, and most relevant to the present work, was the use of activity maps that were not initially designed to measure motor or cognitive habits, but rather, general human behaviour. Hence, one strength of the current work is the study of common hubs for different measures not initially intended to be directly compared. Ultimately, some contrasts are based upon validated mechanisms part of habitual behaviour, such as experimental situations (stress or time pressure) known to largely engage habits and were therefore selected from certain studies;

### Data analysis

The latest version of the GingerALE software v3.0.2 (Eickhoff et al., 2009) was used to compute activation likelihood estimations (ALE) (Turkeltaub et al., 2002). From each of the accepted articles, the coordinates of peak activations were manually extracted and those in Talairach space were transformed to Montreal Neurological Institute (MNI) space using the inverse transform of icbm2tal from GingerALE (Lancaster et al., 2007). The ALE method was implemented as follows: (1) for each study a 3D Gaussian distribution was created around every peak activation (variance proportional to the sample size of the study). This allowed us to use the higher statistical power to reduce peak uncertainty in studies with larger sample sizes. This step was done for every peak to produce one modelled activation (MA) map per study. The value for each voxel in a MA-map represented the probability of that voxel containing an activation foci. (2) Voxel probabilities in all MA-maps were merged to produce an ALE map. Following recommendations of Eickhoff et al., (2016), the ALE map was thresholded using a cluster-level Family-Wise Error (FWE), with a cluster-forming threshold of .001 and a cluster-level FWE of .05 (as stated in the GingerALE manual; Fox et al., 2013).

The first step in applying a cluster-level FWE was to threshold the ALE map at the voxel-level (cluster-forming threshold). To do this, all possible combinations between all peaks from each MA-map were tested. The ALE values provided a null distribution which assumed the peaks would be randomly distributed following all potential combinations amongst them. Here, an uncorrected *p*-value was used since in the next step, a FWE correction was applied to correct for the possibility of multiple comparison errors.

Then, an additional threshold (cluster-level FWE) was applied to select only the largest clusters. Peaks in every MA-map were distributed randomly and then combined into a single ALE map, following the same union method described above for the voxel-level threshold. The ALE maps were also thresholded with the same method used at the voxel-level threshold. This procedure was repeated (1000 times for this meta-analysis) using Monte Carlo permutations method, selecting on every run the largest cluster in the thresholded ALE map (null distribution). Finally, we applied the selected threshold (0.05) to obtain the FWE-corrected results. Using a 0.05 cluster-level threshold resulted in a thresholded map where only 5% of the surviving clusters could have been introduced by chance (false positives), following guidelines to discard non-significant clusters (Eickhoff et al., 2009).

Contrasts between thresholded ALE maps were computed to compare between our conditions of interest (e.g. Everyday-life > Experimental maps). All foci in the selected studies were pooled into a single dataset, which we then split in two datasets by randomly assigning to each one the same number of foci in their original file. The ALE-scores of these two random maps were then subtracted in a voxel-wise manner generating the map of their ALE-scores differences (repeated 10000 times to record ALE-scores and generate a null distribution of randomly spatially distributed foci). Finally, the actual two ALE maps corresponding to the contrast were combined voxel-wise by subtracting the ALE-score of one map from the other. The resulting map of ALE-scores were then thresholded voxel-by-voxel by comparing them with the null distribution previously obtained. We used the default *p*-value suggested in GingerALE (0.01 uncorrected). All computations were run via in-house python scripts to automate analyses in the GingerALE’s interface. Thresholded maps are reported in MNI152 space (Mazziotta et al., 2001).

### Complementary analysis

To ensure the results were not driven by a small number of studies from specific categories (i.e. heterogeneity), several additional tests were conducted (as indicated by Müller et al., 2018). We categorized possible differences by dividing each category (experimental, everyday-life) into subgroups and analysed independently. Studies of experimental habits were separated into two subcategories: probabilistic learning vs other experimental tasks (Table S2), on the ground that probabilistic classification typically activates more anterior portions of the caudate and the putamen, compared with other tasks (Patterson and Knowlton, 2018). Meanwhile, we included categorical variables that separated motor (walking, driving) and cognitive (reading, writing) habits acquired in everyday-life, a further sub-division motivated by the anatomo-functional gradient along fronto-striatal circuits (Haynes and Haber, 2013). To best isolate the sensorimotor properties of habitual driving as part of the motor category, we selected contrasts including less cognitive demands. We also ensured to include > 20 studies in the experimental and everyday-life categories in order to fulfil the expected effect (Eickhoff et al., 2016). Finally, we also performed an additional check for heterogeneity and individual study bias which consists in counting the number of studies that contribute to a specific ROI which was significant in the ALE analysis (Müller et al., 2018). To do so, we applied an FWHM smoothing, whose value depended on the number of subjects per study which was configured according to recommendations in the GingerALE manual. Then, we reviewed the clusters to check if the foci of each study had any influence for each ROI that resulted significant in the group analysis.

## Results

### Study selection

First, to select a cohort of studies investigating everyday-life habits, we analysed data from 57 studies (a total of 1536 subjects) that used diverse stimulus-response paradigms (walking, reading, writing and driving). Imaging models that included automatic parameters on each task were chosen (Tables S1-S2). Second, we sought to confirm the neural basis of newly learned experimental habits in the 38 studies (a total of 938 subjects) that had used probabilistic or discriminative learning, 2-step learning or sequential tasks (Box 1B).

### Striatal role in everyday-life and experimental habits

To find the neural signatures of everyday-life habits, we obtained cluster activity associated with stimulus-response behaviours learned throughout life, and compared these with the signatures of habits newly learnt in laboratory settings. The main effects of naturalistic habits revealed significant bilateral activity in the posterior putamen (Table 1; Figure 2A). Specifically, this was sustained by activity in dorsal sections and left putamen activation that expanded to its posterior boundary (Table 1; Figure 2A-B). For these tasks, other active regions were seen in the cerebellum and cortical areas including the premotor, SMA and inferior frontal gyrus pars opercularis (Table 1; Figure 2A-B).

**Figure 2.**
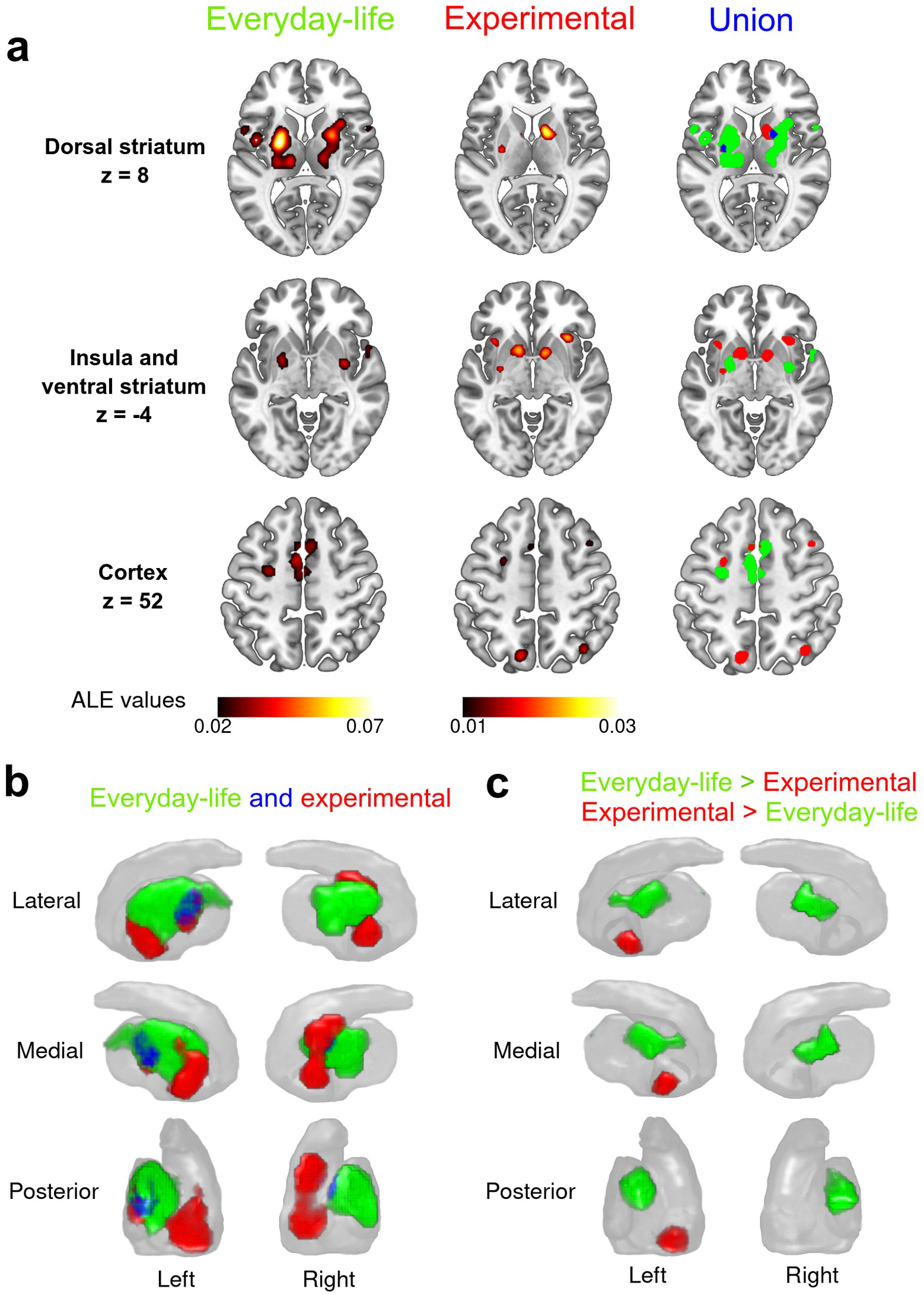
Everyday-life and experimental thresholded activation maps. **a)** Axial views for the main regions. Overlap regions at the right column are shown in blue. Note that there exists activation in the cerebellar cortex for the case of everyday-life studies but it is omitted here for brevity. Z=52 view for the experimental studies is shown as an unthresholded map for visualization purposes (ALE value ≈ 0.01). **b)** 3D striatum reconstruction showing all the activation that fall inside it. Overlap regions at the right column are shown in blue. **c)** 3D striatum reconstruction showing the differential activation of the Everyday -life > Experimental contrast (in green) and the Experimental > Everyday-life contrast (in red).

**Table 1.** Everyday-life habits peaks coordinates in MNI152 space with region names from Harvard-Oxford atlas. Percentages of each brain region indicate how much activation from a cluster fall into such region. Coordinates of any activation comprising < 5% of its volume in a region are not shown for conciseness.

| Cluster ID | Volume (mm<sup>3</sup>) | Brain regions (%) | MNI peak coordinates |  |  | ALE value |
| --- | --- | --- | --- | --- | --- | --- |
|  |  |  | x | y | z |  |
| 1 | 9232 | Left Putamen (54 %) | -24 | -4 | 6 | 0.078 |
|  |  | Left Thalamus (28 %) | -14 | -22 | 10 | 0.04 |
|  |  | Left Pallidum (16 %) | -23 | -4 | 2 | 0.058 |
| 2 | 6616 | Right Putamen (57 %) | 24 | 4 | 6 | 0.056 |
|  |  | Right Thalamus (18 %) | 16 | -16 | 8 | 0.035 |
|  |  | Right Insula (12 %) | 32 | 14 | 8 | 0.037 |
|  |  | Right Pallidum (11 %) | 22 | -8 | 8 | 0.04 |
| 3 | 4840 | Left Supplementary Motor Area (40 %) | -6 | 0 | 52 | 0.035 |
|  |  | Right Paracingulate Gyrus (16 %) | 6 | 14 | 50 | 0.031 |
|  |  | Left Paracingulate Gyrus (16 %) | 0 | 8 | 53 | 0.031 |
|  |  | Right Supplementary Motor Area (14 %) | 4 | 8 | 56 | 0.03 |
| 4 | 3088 | Left Precentral Gyrus (97 %) | -54 | -2 | 40 | 0.038 |
| 5 | 2288 | Right-Cerebellum-Cortex (100 %) | 8 | -64 | -20 | 0.037 |
| 6 | 2288 | Left Central Opercular Cortex (54 %) | -46 | -2 | 6 | 0.037 |
|  |  | Left Inferior Frontal Gyrus pars opercularis (19 %) | -54 | 8 | 24 | 0.027 |
|  |  | Left Precentral Gyrus (17 %) | -56 | 8 | 4 | 0.033 |
|  |  | Left Insula (7 %) | -44 | -1 | 4 | 0.03 |
| 7 | 2152 | Right Precentral Gyrus (76 %) | 56 | 12 | 32 | 0.041 |
|  |  | Right Inferior Frontal Gyrus pars opercularis (19 %) | 54 | 13 | 31 | 0.038 |
| 8 | 1072 | Right Inferior Frontal Gyrus pars opercularis (34 %) | 58 | 12 | 10 | 0.027 |
|  |  | Right Central Opercular Cortex (27%) | 52 | 6 | 2 | 0.029 |
|  |  | Right Temporal Pole (13 %) | 54 | 10 | -2 | 0.034 |
|  |  | Right Precentral Gyrus (12 %) | 3 | -10 | 53 | 0.022 |
|  |  | Right Planum Polare (9 %) | 53 | 5 | -3 | 0.027 |
| 9 | 1016 | Left Precentral Gyrus (43 %) | -26 | -8 | 56 | 0.04 |
|  |  | Left Superior Frontal Gyrus (35 %) | -4 | 14 | 53 | 0.029 |
|  |  | Left Middle Frontal Gyrus (21 %) | -51 | 4 | 38 | 0.027 |

Newly learned habits from laboratory settings were also associated with increased levels of activation in putaminal sections, but with larger representations in anterior striatum (Figure 2C). A gradient was seen along the rostro-ventral section of the putamen, right caudate and the nucleus accumbens bilaterally (Table 2; Figure 2C). The right insula was one of the extrastriatal hubs of activity associated with habits acquired in the laboratory (Table 2; Figure 2A).

**Table 2.** Experimental habits peaks coordinates in MNI152 space with region names from Harvard-Oxford atlas. Percentages of each brain region indicate how much activation from a cluster fall into such region. Coordinates of any activation comprising < 5% of its volume in a region are not shown for conciseness.

| Cluster ID | Volume (mm<sup>3</sup>) | Brain regions (%) | MNI peak coordinates |  |  | ALE value |
| --- | --- | --- | --- | --- | --- | --- |
|  |  |  | x | y | z |  |
| 1 | 3000 | Right Caudate (48 %) | 12 | 6 | 10 | 0.028 |
|  |  | Right Pallidum (23 %) | 14 | 4 | 0 | 0.023 |
|  |  | Right Accumbens (12 %) | 10 | 8 | -6 | 0.03 |
|  |  | Right Putamen (8 %) | 14 | 9 | -6 | 0.02 |
|  |  | Right Thalamus (7 %) | 12 | -1 | 10 | 0.018 |
| 2 | 2136 | Left Putamen (30 %) | -14 | 5 | -9 | 0.027 |
|  |  | Left Accumbens (28 %) | -12 | 6 | -10 | 0.033 |
|  |  | Left Caudate (21 %) | -10 | 4 | 4 | 0.017 |
| 3 | 784 | Left Putamen (99 %) | -28 | -10 | 6 | 0.02 |
| 4 | 752 | Right Insula (90 %) | 32 | 22 | -4 | 0.023 |
|  |  | Right Orbito-Frontal Cortex (10 %) | 34 | 22 | -7 | 0.016 |
| 5 | 736 | Left Insula (90 %) | -34 | 16 | 2 | 0.025 |
|  |  | Left Frontal-Operculum Cortex (10 %) | -36 | 19 | 2 | 0.02 |

Of particular interest was the possibility that despite the differences between neural activation patterns associated with everyday and experimental habits, there may be a common neural substrate accessed by both categories of stimulus-response behaviour. Specifically, we found common activations in anterior right putamen and posterior left putamen (Figure 2A). Although both categories of habits recruited dorsal sub-regions of the putamen bilaterally, the activation by everyday-life habits was stronger (Figure 2C). As expected, no activation in the caudate nucleus survived FWE-corrected thresholds with everyday-life habits. In contrast, habits acquired in the laboratory showed a differential recruitment of the nucleus accumbens, and to a lesser extent, the most antero-ventral section of the right caudate nucleus and putamen (Figure 2C). Hence, while the striatum is a common hub for both categories of habitual behaviour, antero-posterior differences were reflected in the activation patterns of habits acquired in everyday-life and the experimental laboratory.

### Motor and associative segregation in life-long habits

We next tested whether the neural correlates of everyday-life habits varied between sensorimotor routines (walking or driving) and cognitive-associative ones (reading or writing). Clustered activation in motor tasks revealed a significant presence along the dorsal putamen compared with the associative tasks (Figure 3A). In addition, motor tasks extended the pattern of activation into more posterior sections of the right putamen, but only dorsally for the left putamen. This cluster was present across all the antero-posterior axes (Figure 3B).

**Figure 3.**
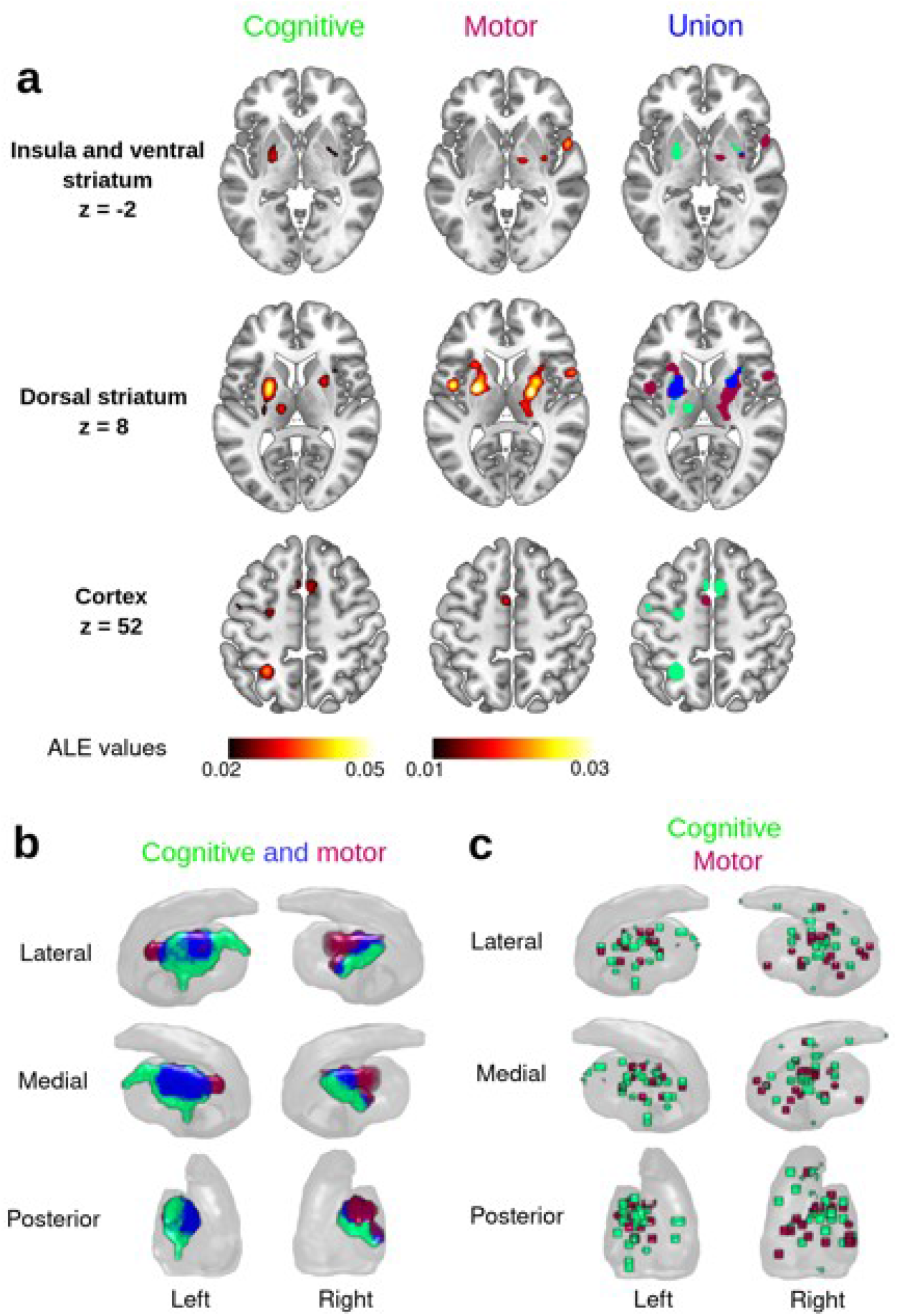
Everyday-life cognitive-motor subcategories thresholded activation maps. **a)** Axial views for the main regions. Overlap regions at the right column are shown in blue. **b)** 3D striatum reconstruction showing all the activation that falls inside it. Overlap regions at the right column are shown in blue. **c)** Foci distribution of all the studies that falls in the striatum. Note that some foci may correspond to the same study.

In the case of habits acquired in everyday-life, we observed an activation pattern in ventral sections of the putamen bilaterally (Table 1). This effect showed significant bilateral asymmetry favoring the left putamen where activation extended into more posterior regions. This was absent in the walking and driving tasks (Figure 3A).

Outside the striatum at the cortical level, the associative tasks acquired in everyday-life recruited premotor area and SMA. Activation was also observed in the cerebellum (Figure 3A). The same cortical regions were recruited during the walking and driving tasks, but none survived FWE-corrected thresholds.

To investigate further the failure of our thresholding procedures to detect activation in the caudate nucleus for any of the naturalistic habit categories, we included the distribution of all the foci that fell into the striatum in both categories of everyday-life habits (Figure 3C). This allowed activity in the right caudate to be observed (Figure 3C) in some associative studies, but not for the motor ones. For the left caudate nucleus, no foci were reported in any study, which confirms the absence of activity reported in the thresholded maps above.

### Probabilistic learning and other tasks segregation in experimental habits

Finally, we aimed to replicate previous findings from laboratory conditions that linked the striatum to the learning and execution of new habits. Consistent striatal activity has been reported when evaluating probabilistic or discriminative learning, 2-step learning and sequential tasks (Patterson and Knowlton, 2018). Here, the intention was to confirm and update these results with more recent findings by separating the studies involving trial-and-error probabilistic reward learning from those using different methodologies for stimulus learning. As predicted, both probabilistic learning and the other tasks showed common regions of striatal activation, with largest clusters in the nucleus accumbens and rostro-ventral sections of the caudate and putamen (Figure 4A). However, only the probabilistic tasks were associated with bilateral recruitment of the most anterior region of the putamen (Figure 4B). These results confirm the previous findings reported by Patterson and Knowlton, (2018). Similarly, the other experimental tasks differed with respect to the probabilistic ones by demonstrating unilateral recruitment of the left rostral caudate nucleus (Figure 4B). Outside the striatum, left insular cortex was activated in other tasks but not by the probabilistic ones (Figure 4A).

**Figure 4.**
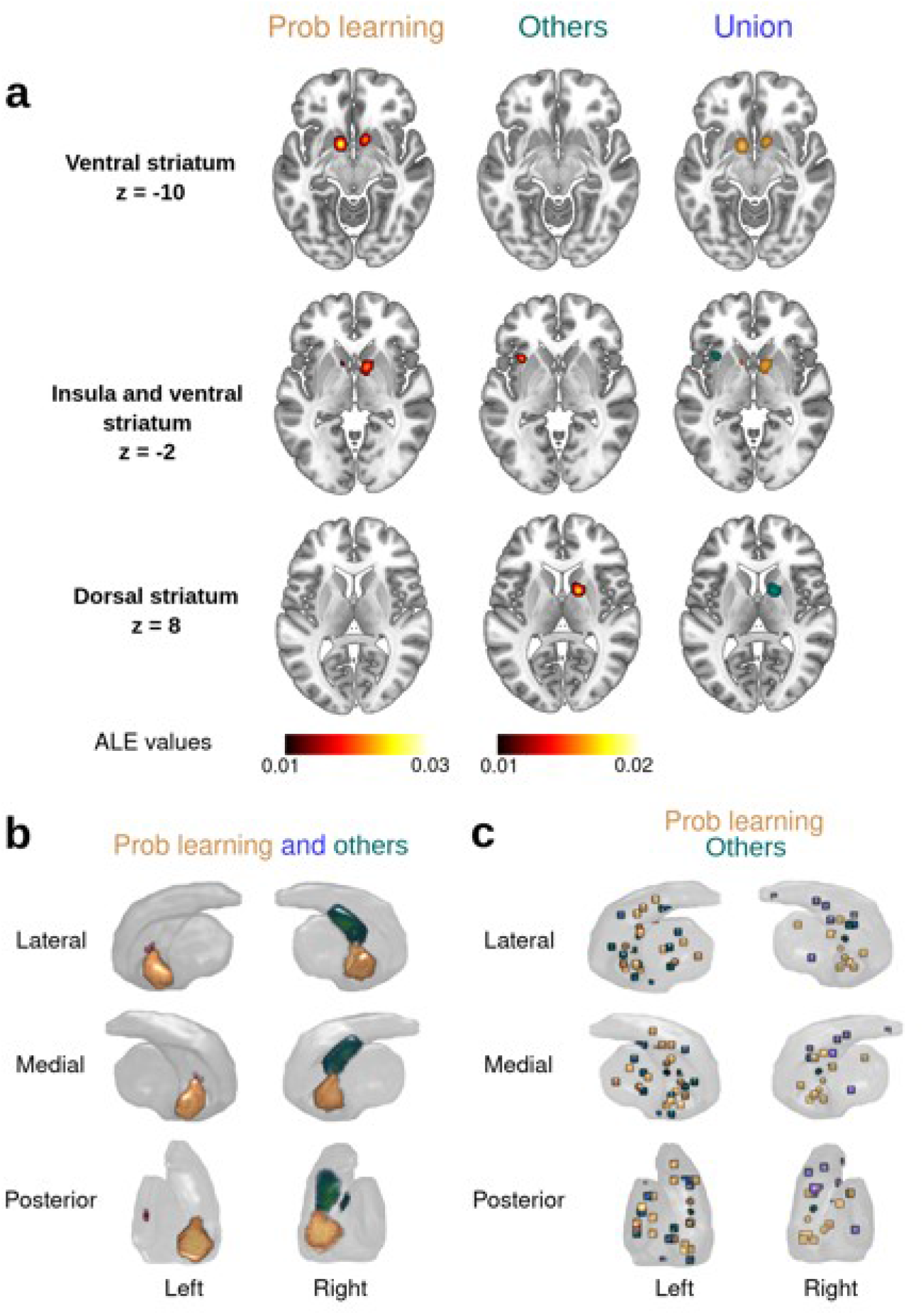
Experimental subcategories thresholded activation maps. **a)** Axial views for the main regions. In this case there are no overlap regions at the right column. **b)** 3D striatum reconstruction showing all the activation that fall inside it. In this case there are no overlap regions at the right column. **c)** Foci distribution of all the studies that fall in the striatum. Note that some foci may correspond to the same study.

### Between-study heterogeneity

The degree of heterogeneity was not strong (that is, a result not based on a reduced number of studies), as was also seen in the subgroup analyses for everyday-life and experimental results (Tables S3-S4).

## Discussion

By comparing long-established habits acquired in everyday-life with newly learned habits acquired under laboratory-controlled conditions, our meta-analysis identified two distinct types of habit-related mechanisms in the human striatum. Both common and distinct functional links between different sub-regions of the brain’s habit-associated circuitry were identified. Differences in neural substrate were found with life-time habits showing enhanced activity in the dorsal posterior putamen, together with activity in the cerebellum and SMA, while laboratory habits engaged anterior sections of the putamen with activation extending into caudate nucleus and nucleus accumbens. Importantly, common regions of activation were found in posterior left and anterior right putamen. Ultimately, delineation of specific striatal contributions to motor-associative variables embedded in habits were responsible for shared anatomical patterns in everyday-life associative habits, such as reading and writing, that engaged rostral regions of the putamen.

### The putamen function in everyday-life habits

The current findings provide direct evidence for bilateral putamen engagement in habits, long acquired and executed in everyday-life. Typically, we observed larger bilateral activity in studies of life-time habits compared with newly acquired experimental ones. This may implicate a broad putaminal role for the stimuli-rich sensorimotor computations required during complex everyday stimulus-response tasks. Consistent with this view, neuroimaging, lesioning and animal electrophysiological investigations all converge to pinpoint a role for the putamen in the expression of associations between specific stimuli and movement patterns to produce habitual behaviours with predicted outcomes (Bellebaum et al., 2008; Cromwell and Schultz, 2003; Haruno and Kawato, 2006; Sommer and Pollmann, 2016; Yamada et al., 2004). Indeed, the neurophysiological properties of the putamen are supported by subpopulations of neurons that respond to sensory stimuli (Gardiner and Nelson, 1992), unifying actions sets for movement sequences (Cellini, 2017; Wymbs et al., 2012) and the integration of elementary units such as individual finger movements (Andersen et al., 2020). Moreover, putaminal activity is not solely dedicated to movement parameters, but is also present while learning without motor plans (Romero et al., 2008), increasing neural response magnitude to reinforced choices (Cohen et al., 2021), and for predicted well-learned and contextually-driven actions (Kimura, 1986; Kimura et al., 1990, 1984; Kunimatsu et al., 2019; Peters et al., 2021; Tunik et al., 2009). Hence, the diversity of sensorimotor-related and high-order reinforcement neurons in the putamen support it having a pivotal role at several levels in context-rich scenarios (sensorimotor control, predicted actions, object values) in the acquisition and expression of stimulus-response habits.

The role of the putamen in habitual behaviour is directly influenced by ascending dopaminergic system acting on cortical and thalamic inputs to this region (Haber et al., 2000). Afferent dopamine signals provide critical modulatory influences on striatal subregions whereby projections from substantia nigra pars compacta and the ventral tegmental area differentially target terminals in dorsal and ventral striatum respectively (Björklund and Dunnett, 2007). The main nigrostriatal projection is topographically organised with a medial to lateral gradient (Matsuda et al., 2009). Tonic firing within this system sustains, motivational, cognitive and action-specific decision making (Hikosaka et al., 2000; Redgrave et al., 1999), while sensory-evoked phasic patterns of dopaminergic activity provide a general mechanism for reinforcement learning associated with the acquisition of novel actions (Redgrave et al., 2008; Schultz, 2006). The large bilateral activity found in the striatum for the routines of everyday-life should be influenced by ascending dopaminergic fibers facilitating the selection of appropriate low-level stimulus-response associations required for writing or walking (proprioceptive and muscle sensorimotor control). Of interest, the dorso-lateral caudal putamen was active for both cognitive (reading, writing) and motor operations (driving, walking), which corresponds with our understanding of parallel inter-related cortico-striatal functions (McFarland and Haber, 2002; Morris and Stein, 2017). The stimulus-response selection role of the putamen may well represent the low level sensorimotor selections necessary in all forms of habitual motor behaviour (Bromberg-Martin et al., 2010; Romero et al., 2008). Hence, the putamen will likely respond preferentially to the predictable sensorimotor contingences present in seemingly sophisticated behaviours that rely, in part, on low-level automatic stimulus-response associations for their execution, driving for example.

### Cortical control over everyday-life habits

The substantial input that the striatum receives from cortical areas (Peters et al., 2021) permits this region to receive contextual, motivational and/or motor cues relevant to action selection (Haber and Knutson, 2009; Redgrave et al., 1999). Several cortical areas represent potentially competing behavioural options that can be expressed under habitual control, including ventral premotor (de Wit et al., 2012b; Ohbayashi et al., 2016), extraestriate visual cortex (Luque et al., 2017) and bilateral insula or precentral gyrus (Eryilmaz et al., 2017). In addition, premotor cortex has been shown to interact directly with the posterior putamen during the acquisition of habits (de Wit et al., 2012b).

In the present study we found that SMA activity was an essential component of the habitual circuitry engaged by the everyday-life tasks. The SMA represents one of the important junctions between cortical-subcortical motor and cognitive circuits (Nachev et al., 2008), with projections to the dorso-lateral striatum and posterior putamen (Leh et al., 2007). The SMA has been shown to be involved in learning stimulus-response contingencies (Chen and Wise, 1995; Nachev et al., 2008) and model-free (habitual) tasks (Morris et al., 2016). Yet, the SMA is not represented in most studies involving new learning of actions. Probably, the multi-dimensional components required while learning to walk or drive (somatosensory, visuo-spatial, prediction, motor planning, etc) will be mediated via other cortical inputs (likely SMA integration function) to the putamen (Haber and Knutson, 2009; Leisman et al., 2016). When bolted together, serial selections in such complicated but eventually predictable tasks can be viewed as coherent sequences of habitual behaviour. For example, the sophisticated set of movements required to change gear when engine revs go above a certain level, and the precise flow of hand movements required to write individual letters, with repetition, can be expressed under automatic stimulus-response control. The absence of SMA activity in experimental tasks included in the present, and previous investigations, attests to the importance of multi-dimensional computations when cognitive and motor functions are integrated to develop habitual sequences of behaviour in everyday-life. Such automatic sequences are likely to recruit medial cortical motor areas including the SMA as a mechanism to boost expected motor and associative computations that, with subcortical structures, including putamen (Cunnington et al., 2002; Smittenaar et al., 2013) represent a looped architecture that can select an appropriate sequences of sensory-driven movements that implement predictable task components.

Our results also showed a consistent activation of the inferior frontal gyrus. but only by everyday-life habits. It is known that this region is part of the mirror neuron system related to action observation and execution (Kilner et al., 2009; Molenberghs et al., 2012). A possible interpretation of this activation would be that mirror neurons encode behaviour at the level of actions, rather than as individual movements comprising complex behaviour. Consequently, only when an action representation is established and well-practiced would a mirror system be expected to respond accordingly. Perhaps recent, laboratory-acquired stimulus-response associations are not sufficiently well embedded.

### Everyday-life vs experimental scenarios to study habits

Our analyses may also have implications for understanding previous studies that investigated novel stimulus-response habits learned in experimental laboratories (probabilistic or discriminative learning, 2-step learning or sequential tasks), and which also report significant striatal activations (de Wit et al., 2012b; Eryilmaz et al., 2017; Huang et al., 2020; Jankowski et al., 2009; Lehéricy et al., 2005b; Liljeholm et al., 2015a; Morris and Stein, 2017; Patterson and Knowlton, 2018; Tricomi et al., 2009). Specifically, our analyses show stimulus-responses behaviour acquired both in everyday-life and under laboratory conditions showed common striatal activity in the right anterior putamen and left posterior putamen. The everyday-life tasks included long established sensorimotor responses triggered by sensory cues in the absence of new associative learning. In contrast, laboratory learned habits typically involved relatively minor motor components (finger key presses), but novel cue-response associations driven by reinforcement learning. In line with a previous meta-analysis on basal ganglia activation across multiple motor disciplines (Arsalidou et al., 2013), left lateralized putaminal activity was prominent in motor operations (such as eye movements and body motion), has a larger volume in right-handed participants (Peterson et al., 1993), and is critical in behaviours guided by stimulus-response mappings (Liljeholm et al., 2015a; Patterson and Knowlton, 2018). Hence, our observation of left posterior putamen activation by both long-established and novel habits suggests this region is critical for aspects of automatic stimulus-response processing that are present in a wide range of habitual behaviours.

In contrast, activation of the anterior putamen would accord well with neural patterns associated with initial learning of stimulus-response associations in experimental settings (Jankowski et al., 2009). Activity in key regions of the circuitry associated with goal-directed behavioural control (caudate and nucleus accumbens) were also present in laboratory studies of habits. These findings match those of several fMRI studies using various reinforcement learning tasks. When encoding the value of predicted reward outcome linked to new actions activation of the ventro-medial prefrontal cortex, insula and anterior striatum has been widely reported (Grabenhorst and Rolls, 2011; Parkes et al., 2015; Tanaka et al., 2008). An issue here could be that in some of these studies behaviours established in the laboratory that are presumed to be habits fail to meet formal requirements. For example, several recent human experimental studies failed to demonstrate an effect of training duration on insensitivity to outcome-devaluation test (de Wit et al., 2018; Pool et al., n.d.; Watson and de Wit, 2018). In such studies it is possible that more trials may be needed to establish stimulus-response associations that are fully automatic and can persist after devalued outcome challenges. In the case of every-day habits, independence of outcome value is typically demonstrated in cases of action slips. That is, when a long established habitual response produces an undesired outcome, such as taking the elevator to the floor of an old office. Last, there is the possibility that some differences between everyday-life and laboratory-developed habits, such as activation of the nucleus accumbens, may be due to the explicit rewards component present in some laboratory paradigms rather than any fundamental difference in the habitual network.

### Dysfunctional habits: inability to exploit stimulus-response habits

Insofar as the caudal putamen has been identified as a critical node in the neural substrate responsible for automatic habitual behaviour, malfunctioning of this region should be associated with deficits in habitual performance. A notable instance of this would be the differential loss of dopamine neurotransmission from this region in Parkinson’s disease (Redgrave et al., 2010a). The causes of this selective loss is not known, however given the high metabolic cost of dopaminergic neurotransmission Hernandez et al., (2019) suggested it could reflect an over-reliance throughout life on automatic habitual control. The particular problems Parkinson’s disease patients have with stimulus-response behaviours e.g. walking and writing (Bannard et al., 2019; Peterson and Horak, 2016; Wu et al., 2016), and their inability to demonstrate novel associations in the laboratory (de Wit et al., 2010; Wu et al., 2015), can be seen as an inability of Parkinson’s patients to enact established habits and acquire new ones. In contrast, other neuropsychiatric conditions, such as the addictions, patients exhibit an excessive cue-dependent use of certain rewards linked to increased posterior putamen activity (Everitt et al., 2008; Sjoerds et al., 2013). In the light of this work, an important future question would be whether the limbic and associative territories of the basal ganglia (Alexander and Crutcher, 1990) can operate in a similar habitual mode as has been established for stimulus-response motor habits in the sensorimotor territories. If so, this would move towards a reconceptualization of stimulus evoked motivations (e.g. drug craving), stimulus-evoked cognitions (e.g. prejudices), and stimulus-evoked motor responses (motor habits) as part of a general mode of model-free stimulus-response behavioural control. Were this to be the case, one might speculate there would likely be a general mechanism of neural implementation across all territories of the basal ganglia.

### Limitations & strengths

Despite the clear positive findings of the present investigation, certain limitations must be acknowledged. First, the ALE analysis can cause some parameters, such as voxel peaks, to be overlooked. Ideally, exploring the statistical activity maps of each study individually would be of great value. Unfortunately, most studies did not have full imaging datasets so this would be precluded.

Second, some studies using RPE contrasts combined mixed regressors of choice and outcome thereby rendering it difficult to make separations between habitual stimulus-response components. To avoid this, we explored our principal findings without these studies and found similar patterns of striatal activation (anterior putamen; Figure 4A). Moreover, our heterogeneity analysis showed a reduced number of RPE studies (6/25 studies for left putamen; 4/20 studies for right putamen) that contributed to the significant activity of such clusters (as displayed in Table S4 for experimental studies). Consequently, the putamen clusters we report are not mainly driven by the RPE studies

Third, as stimulus-response habits are driven by specific conjunctions of sensory stimuli, the heterogeneity of every-day and laboratory tasks included in our analysis means the stimuli driving brain responses necessarily will have differed. Moreover, the laboratory studies included have minor motor components compared with the typical motor-sophistication of habits established over a lifetime. To control for this possibility, and following imaging meta-analysis best practice (Müller et al., 2018), as different stimuli and task-related activities were collected, general and specific sub-analysis were conducted. Last, due to the difficulty of performing some behaviours in the MRI scanner, necessarily we had to restrict our selection to everyday-life behaviours where scanning was possible. This could restrict the generalisation of our findings to other activities prevalent in every-day life.

The main strengths of the work point to the possible generalizability and external validity of lab-developed habits, as well as proposing novel methodological procedures to investigate human habitual behaviour. The heterogeneity of tasks necessarily means there was a diverse range of both stimuli and behavioural responses deployed in the studies included in our analyses. It is therefore remarkable we were able to pull out common patterns of neural activation associated with stimulus-response control.

A further strength of our investigation was that we were able to find significant striatal activation in tasks with high levels of stimulus-response behaviour in studies that were not originally designed to study habits. This was despite the behaviours of interest not having been formally identified as habitual using outcome devaluation or other tests. That we find common neural maps with widely different means for testing habits lends support to this idea.

### Future research and conclusions

Recently, a range of procedures have been developed to establish and test the expression of newly acquired habits in the experimental laboratory, including reversal of learned actions after overtraining (Luque et al., 2020), overloading goal-directed top-down control while measuring execution of learned stimulus-response associations (Hardwick et al., 2019) or biasing movement kinematics (Wong et al., 2017). Interestingly, pre-existing categorical associations established in everyday-life (i.e. color associations or prejudice) have a clear advantage on measuring automatic processing (Amodio, 2014; Ceceli et al., 2020). Further methodological options for studying long established habits acquired in everyday-life include assessing the motor performance of expert musicians (Vaquero et al., 2016), tennis players (Cacioppo et al., 2014) and expert shooters (Kim et al., 2014). Hence, getting subjects to bring their everyday-life stimulus-response associations into the laboratory under controlled conditions is an important option for studying the neural substrates underlying habitual behaviour in humans. Although there is less control of the independent variables in such studies, by selecting subjects with different amounts of everyday-life experience it is possible to relate the amount of practice with habit strength.

In conclusion, the present study points to a fundamental functional role for the posterior putamen in the expression of habits acquired in everyday-life. Importantly, a critical dissociation was established between different brain regions whose contributions to motor-cognitive representations of habits have, thus far, been largely indistinguishable. Conversely, the engagement of the anterior putamen was shown to be associated more with habits newly learned in experimental laboratories. Careful experimental protocols must be designed to identify those chunks of complex behaviour that are under automatic stimulus-response control and can occur independently of outcome value or response contingency. This is true both for newly acquired associations in the laboratory and long-standing habits established in everyday-life. Finally, the present study highlights the importance and value of having subjects bring life-long habits into the laboratory to be investigated and compared with recently acquired stimulus-response associations. This meta-analysis provides strong evidence for a striatal role in everyday-life and lab-developed habits giving a novel contribution to the literature on the general neural footprint of human habits.

## Acknowledgments

We are thankful to Blanca Caijao and Cristina Rubio (Universidad Rey Juan Carlos) for their help in article search. MM is funded by a PhD Fellowship from Instituto de Salud Carlos III (PFIS-FI720/00068). IO is funded by Instituto de Salud Carlos III (Miguel Servet, CP18/00038) and AES-ISCIII (PI19/00298) from Ministry of Science and Innovation in Spain. The funders played no role in the idea, design, data collection or analysis, decision to publish or manuscript editing and writing.

## Author Contributions

P.G. design, data collection, writing; M.M. design, data collection, analysis, writing; P.R. writing; D.L. design, data collection, writing; I.O. design, data collection, writing;

## Supplementary material

### Methods

The complete set of search terms were as follows:

1. Search on PubMed Everyday-life habits: (walking[MeSH Terms] OR walking[Title/Abstract]) OR (Driving behaviour[MeSH Terms] OR Driving behaviour[Title/Abstract]) OR (Driving behavior[MeSH Terms] OR Driving behavior[Title/Abstract]) OR (car driving[MeSH Terms] OR Car driving[Title/Abstract]) OR (writing[MeSH Terms] OR writing[Title/Abstract]) OR (handwriting[MeSH Terms] OR handwriting[Title/Abstract]) OR (reading[Title/Abstract] OR reading[MeSH Terms) AND (functional magnetic resonance imaging[MeSH Terms] OR fMRI[Title/Abstract]) AND ((basal ganglia[Title/Abstract] OR caudate[Title/Abstract] OR putamen[Title/Abstract] OR striatum[Title/Abstract]) OR (basal ganglia[MeSH Terms] OR caudate nucleus[MeSH Terms] OR putamen[MeSH Terms] OR striatum[MeSH Terms])) Experimental habits: (“habit” OR “habits” OR “probabilistic classification” OR “weather prediction” OR “response learning” OR “instrumental conditioning” OR “instrumental learning” OR “reinforcement learning” OR “outcome devaluation” OR “sequential decision” OR “two step” OR “2 step”) AND (“basal ganglia” OR “caudate” OR “putamen” OR “striatum”) AND (“fMRI” OR “functional magnetic resonance imaging” OR “functional MRI”) AND (“2017/06/22”[PDAT] : “3000/12/31”[PDAT])
2. Search on Web of Science Everyday-life habits: TI=(((driving) OR (car driving) OR (walking) OR (writing) OR (handwriting) OR (reading)) AND ((functional magnetic resonance imaging OR fMRI)) AND (((basal ganglia OR caudate OR putamen OR striatum)))) OR AB=(((driving) OR (car driving) OR (walking) OR (writing) OR (handwriting) OR (reading)) AND ((functional magnetic resonance imaging OR fMRI)) AND (((basal ganglia OR caudate OR putamen OR striatum)))) Experimental habits: (TI=((habit OR habits OR probabilistic classification OR weather prediction OR response learning OR instrumental conditioning OR instrumental learning OR reinforcement learning OR outcome devaluation OR sequential decision OR two step OR 2 step) AND (basal ganglia OR caudate OR putamen OR striatum) AND (fMRI OR functional magnetic resonance imaging OR functional MRI))) OR AB=((habit OR habits OR probabilistic classification OR weather prediction OR response learning OR instrumental conditioning OR instrumental learning OR reinforcement learning OR outcome devaluation OR sequential decision OR two step OR 2 step) AND (basal ganglia OR caudate OR putamen OR striatum) AND (fMRI OR functional magnetic resonance imaging OR functional MRI))

### Supplementary Material Figure Legends and Tables

**Figure S1.**
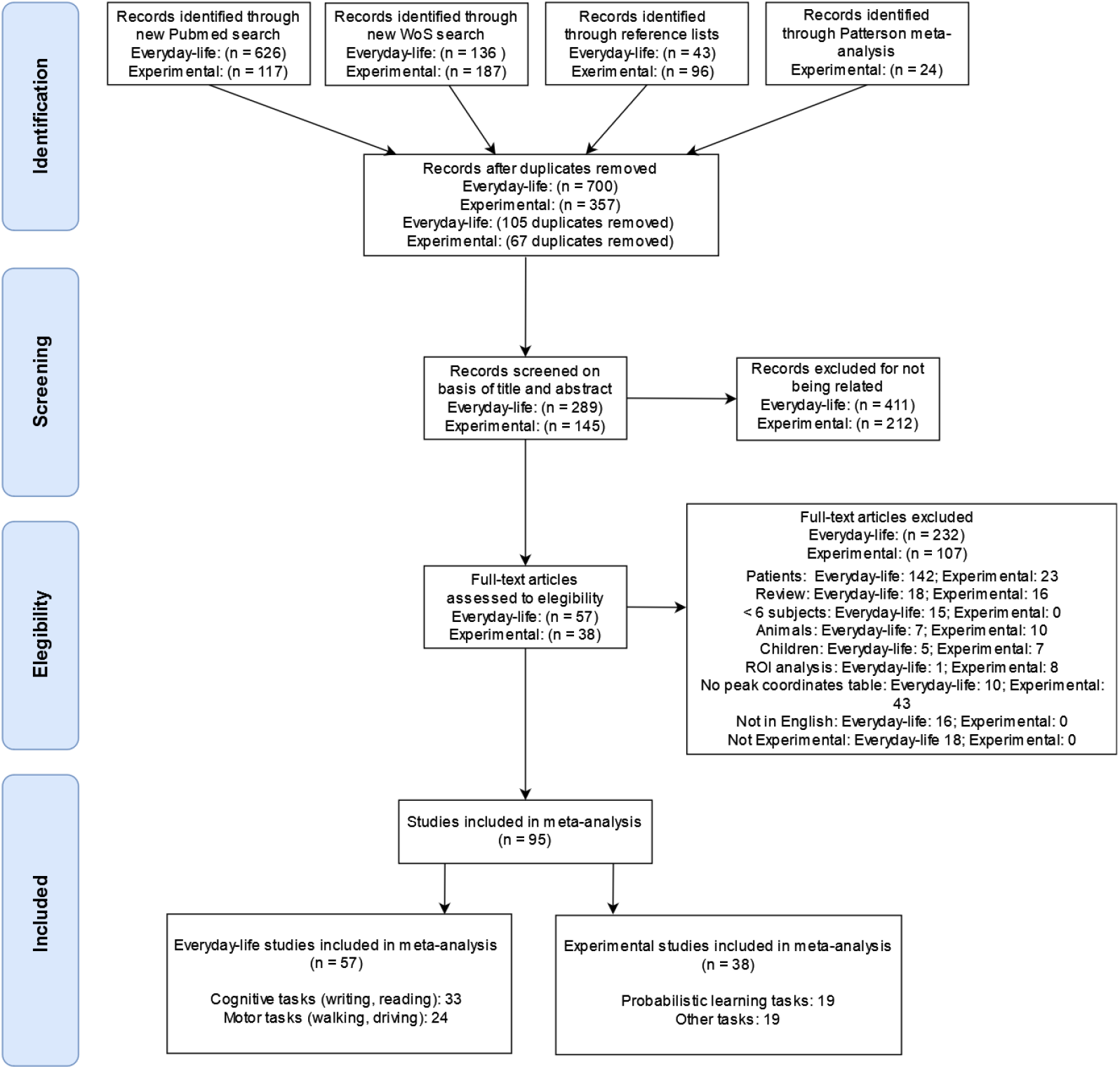
PRISMA chart describing the steps followed for the selection of studies.

**Table S1.**
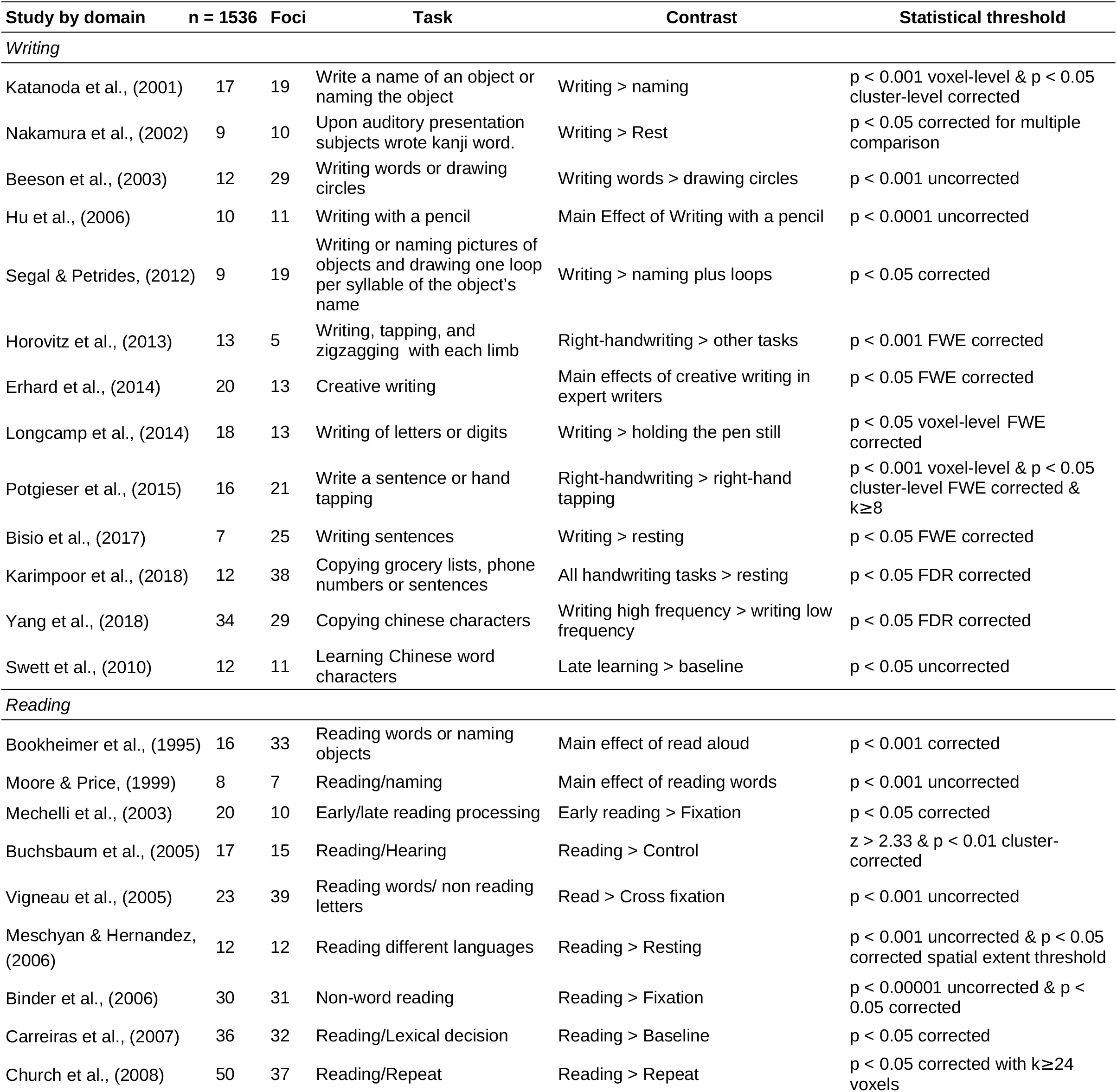
Studies with everyday-life habits included in the meta-analysis.

**Table S2.**
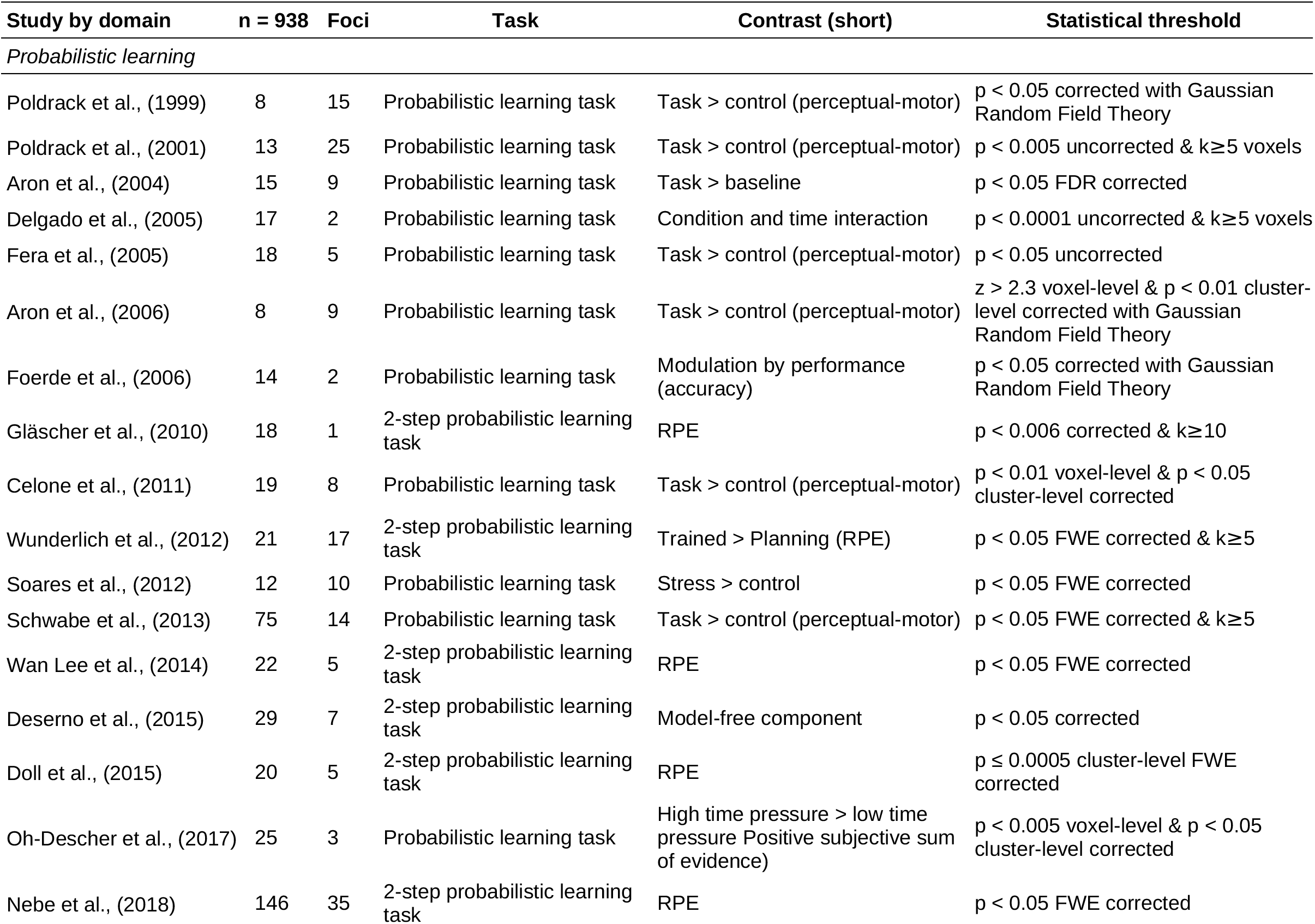
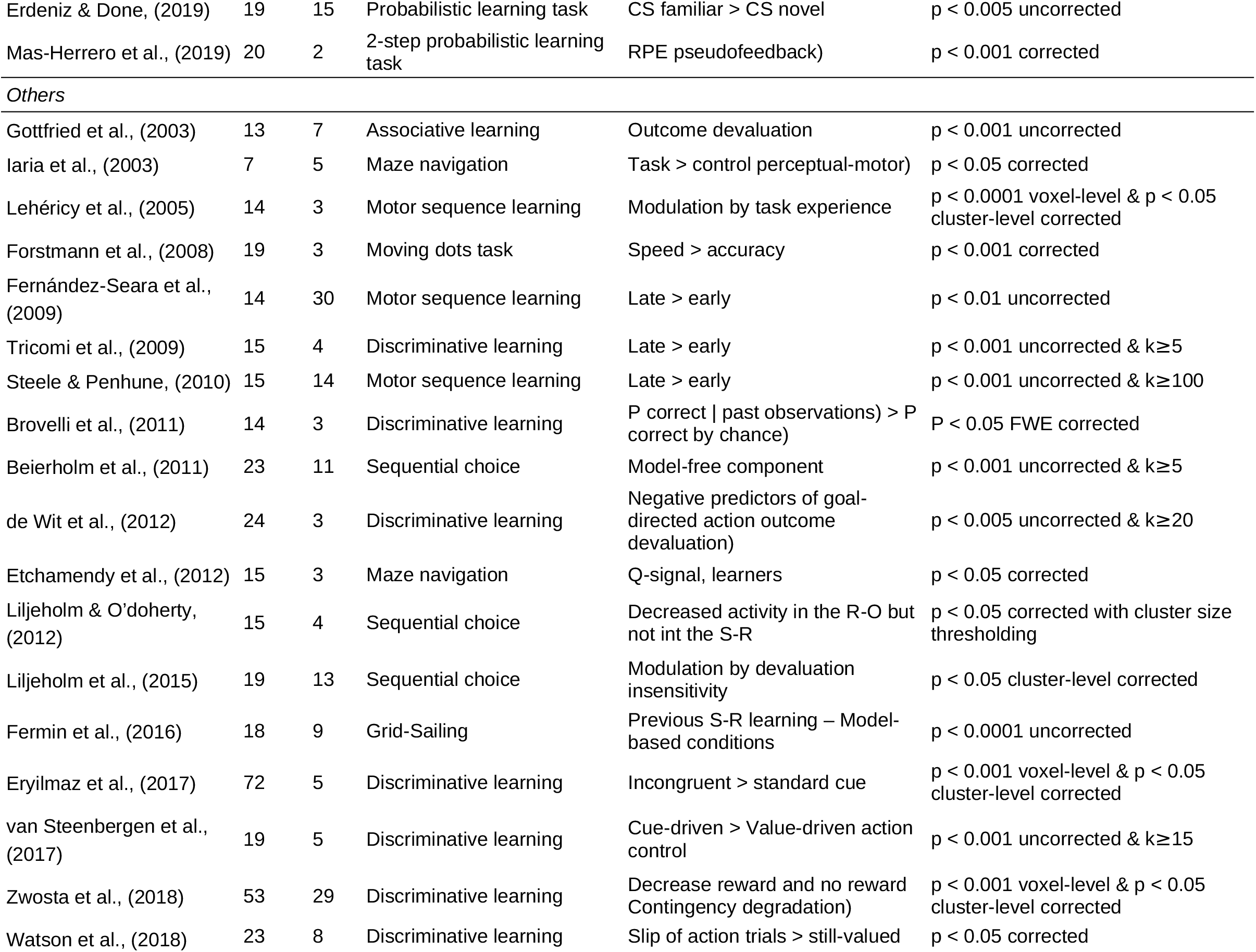
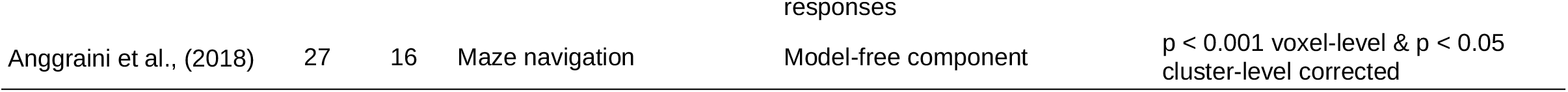
Studies with experimental habits included in the meta-analysis.

**Table S3.** Additional diagnostics used to check the heterogeneity for everyday-life studies. The list details which studies have any foci contributing to the significant clusters in the ALE meta-analysis for everyday-life studies.

(IN CSV)

**Table S4.** Additional diagnostics to check the heterogeneity for experimental studies. The list details which studies have any foci contributing to the significant clusters in the ALE meta-analysis for experimental studies.

(IN CSV)

## References

Adams, C.D., 1982. Variations in the Sensitivity of Instrumental Responding to Reinforcer Devaluation. Q. J. Exp. Psychol. Sect. B 34, 77–98. https://doi.org/10.1080/14640748208400878

Alexander, G.E., Crutcher, M.D., 1990. Functional architecture of basal ganglia circuits: neural substrates of parallel processing. Trends Neurosci. https://doi.org/10.1016/0166-2236(90)90107-L

Allali, G., Montembeault, M., Brambati, S.M., Bherer, L., Blumen, H.M., Launay, C.P., Liu-Ambrose, T., Helbostad, J.L., Verghese, J., Beauchet, O., 2019. Brain structure covariance associated with gait control in aging. Journals Gerontol. -Ser. A Biol. Sci. Med. Sci. 74, 705–713. https://doi.org/10.1093/gerona/gly123

Amodio, D.M., 2014. The neuroscience of prejudice and stereotyping. Nat. Rev. Neurosci. 15, 670–682. https://doi.org/10.1038/nrn3800

Andersen, K.W., Madsen, K.H., Siebner, H.R., 2020. Discrete finger sequences are widely represented in human striatum. Sci. Rep. 10, 1–12. https://doi.org/10.1038/s41598-020-69923-x

Anggraini, D., Glasauer, S., Wunderlich, K., 2018. Neural signatures of reinforcement learning correlate with strategy adoption during spatial navigation. Sci. Reports 2018 81 8, 1–14. https://doi.org/10.1038/s41598-018-28241-z

Aron, A.R., Gluck, M.A., Poldrack, R.A., 2006. Long-term test-retest reliability of functional MRI in a classification learning task. Neuroimage 29, 1000–1006. https://doi.org/10.1016/J.NEUROIMAGE.2005.08.010

Aron, A.R., Shohamy, D., Clark, J., Myers, C., Gluck, M.A., Poldrack, R.A., 2004. Human midbrain sensitivity to cognitive feedback and uncertainty during classification learning. J. Neurophysiol. 92, 1144–1152. https://doi.org/10.1152/JN.01209.2003

Arsalidou, M., Duerden, E.G., Taylor, M.J., 2013. The Centre of the Brain : Topographical Model of Motor, Cognitive, Affective, and Somatosensory Functions of the Basal Ganglia 3054, 3031–3054. https://doi.org/10.1002/hbm.22124

Balleine, B.W., O’Doherty, J.P., 2010. Human and rodent homologies in action control: Corticostriatal determinants of goal-directed and habitual action. Neuropsychopharmacology 35, 48–69. https://doi.org/10.1038/npp.2009.131

Bannard, C., Leriche, M., Bandmann, O., Brown, C.H., Ferracane, E., Sánchez-Ferro, Á., Obeso, J., Redgrave, P., Stafford, T., 2019. Reduced habit-driven errors in Parkinson’s Disease. Sci. Rep. 9, 1–8. https://doi.org/10.1038/s41598-019-39294-z

Beeson, P.M., Rapcsak, S.Z., Plante, E., Chargualaf, J., Chung, A., Johnson, S.C., Trouard, T.P., 2003. The neural substrates of writing: A functional magnetic resonance imaging study. Aphasiology 17, 647–665. https://doi.org/10.1080/02687030344000067

Beierholm, U.R., Anen, C., Quartz, S., Bossaerts, P., 2011. Separate encoding of model-based and model-free valuations in the human brain. Neuroimage 58, 955–962. https://doi.org/10.1016/j.neuroimage.2011.06.071

Bellebaum, C., Koch, B., Schwarz, M., Daum, I., 2008. Focal basal ganglia lesions are associated with impairments in reward-based reversal learning. Brain 131, 829–841. https://doi.org/10.1093/brain/awn011

Binder, J.R., Medler, D.A., Westbury, C.F., Liebenthal, E., Buchanan, L., 2006. Tuning of the human left fusiform gyrus to sublexical orthographic structure. Neuroimage 33, 739– 748. https://doi.org/10.1016/j.neuroimage.2006.06.053

Bisio, A., Pedullà, L., Bonzano, L., Tacchino, A., Brichetto, G., Bove, M., 2017. The kinematics of handwriting movements as expression of cognitive and sensorimotor impairments in people with multiple sclerosis. Sci. Rep. 7, 1–10. https://doi.org/10.1038/s41598-017-18066-7

Björklund, A., Dunnett, S.B., 2007. Dopamine neuron systems in the brain: an update. Trends Neurosci. https://doi.org/10.1016/j.tins.2007.03.006

Bookheimer, S.Y., Zeffiro, T.A., Blaxton, T., Gaillard, W., Theodore, W., 1995. Regional cerebral blood flow during object naming and word reading. Hum. Brain Mapp. 3, 93– 106. https://doi.org/10.1002/hbm.460030206

Bromberg-Martin, E.S., Matsumoto, M., Hikosaka, O., 2010. Dopamine in Motivational Control: Rewarding, Aversive, and Alerting. Neuron. https://doi.org/10.1016/j.neuron.2010.11.022

Brovelli, A., Nazarian, B., Meunier, M., Boussaoud, D., 2011a. Differential roles of caudate nucleus and putamen during instrumental learning. Neuroimage 57, 1580–90. https://doi.org/10.1016/j.neuroimage.2011.05.059

Brovelli, A., Nazarian, B., Meunier, M., Boussaoud, D., 2011b. Differential roles of caudate nucleus and putamen during instrumental learning. Neuroimage 57, 1580–1590. https://doi.org/10.1016/j.neuroimage.2011.05.059

Buchsbaum, B.R., Olsen, R.K., Koch, P.F., Kohn, P., Kippenhan, J.S., Berman, K.F., 2005. Reading, hearing, and the planum temporale. Neuroimage 24, 444–454. https://doi.org/10.1016/j.neuroimage.2004.08.025

Cacioppo, S., Fontang, F., Patel, N., Decety, J., Monteleone, G., Cacioppo, J.T., 2014. Intention understanding over T: A neuroimaging study on shared representations and tennis return predictions. Front. Hum. Neurosci. 8, 781. https://doi.org/10.3389/fnhum.2014.00781

Callan, A.M., Osu, R., Yamagishi, Y., Callan, D.E., Inoue, N., 2009. Neural correlates of resolving uncertainty in driver’s decision making. Hum. Brain Mapp. 30, 2804–2812. https://doi.org/10.1002/hbm.20710

Carreiras, M., Mechelli, A., Estévez, A., Price, C.J., 2007. Brain activation for lexical decision and reading aloud: Two sides of the same coin? J. Cogn. Neurosci. 19, 433–444. https://doi.org/10.1162/jocn.2007.19.3.433

Ceceli, A.O., Myers, C.E., Tricomi, E., 2020. Demonstrating and disrupting well-learned habits. PLoS One 15, 1–28. https://doi.org/10.1371/journal.pone.0234424

Cellini, N., 2017. Memory consolidation in sleep disorders. Sleep Med. Rev. 35, 101–112. https://doi.org/10.1016/j.smrv.2016.09.003

Celone, K.A., Thompson-Brenner, H., Ross, R.S., Pratt, E.M., Stern, C.E., 2011. An fMRI investigation of the fronto-striatal learning system in women who exhibit eating disorder behaviors. Neuroimage 56, 1749–1757. https://doi.org/10.1016/J.NEUROIMAGE.2011.03.026

Cheema, K., Lantz, N., Cummine, J., 2018. Exploring the role of subcortical structures in developmental reading impairments: Evidence for subgroups differentiated by caudate activity. Neuroreport 29, 271–279. https://doi.org/10.1097/WNR.0000000000000938

Chein, J.M., Albert, D., O’Brien, L., Uckert, K., Steinberg, L., 2011. Peers increase adolescent risk taking by enhancing activity in the brain’s reward circuitry. Dev. Sci. 14. https://doi.org/10.1111/j.1467-7687.2010.01035.x

Chen, L.L., Wise, S.P., 1995. Neuronal activity in the supplementary eye field during acquisition of conditional oculomotor associations. J. Neurophysiol. 73, 1101–1121. https://doi.org/10.1152/jn.1995.73.3.1101

Choi, M.H., Kim, H.S., Yoon, H.J., Lee, J.C., Baek, J.H., Choi, J.S., Tack, G.R., Min, B.C., Lim, D.W., Chung, S.C., 2017. Increase in brain activation due to subtasks during driving: FMRI study using new MR-compatible driving simulator. J. Physiol. Anthropol. 36, 1–12. https://doi.org/10.1186/s40101-017-0128-8

Christensen, M.S., Lundbye-Jensen, J., Petersen, N., Geertsen, S.S., Paulson, O.B., Nielsen, J.B., 2007. Watching your foot move - An fMRI study of visuomotor interactions during foot movement. Cereb. Cortex 17, 1906–1917. https://doi.org/10.1093/cercor/bhl101

Chung, S.C., Choi, M.H., Kim, H.S., You, N.R., Hong, S.P., Lee, J.C., Park, S.J., Baek, J.H., Jeong, U.H., You, J.H., Lim, D.W., Kim, H.J., 2014. Effects of distraction task on driving: A functional magnetic resonance imaging study, in: Bio-Medical Materials and Engineering. IOS Press, pp. 2971–2977. https://doi.org/10.3233/BME-141117

Church, J.A., Coalson, R.S., Lugar, H.M., Petersen, S.E., Schlaggar, B.L., 2008. A developmental fMRI study of reading and repetition reveals changes in phonological and visual mechanisms over age. Cereb. Cortex 18, 2054–2065. https://doi.org/10.1093/cercor/bhm228

Ciccarelli, O., Toosy, A.T., Marsden, J.F., Wheeler-Kingshott, C.M., Sahyoun, C., Matthews, P.M., Miller, D.H., Thompson, A.J., 2005. Identifying brain regions for integrative sensorimotor processing with ankle movements. Exp. Brain Res. 166, 31–42. https://doi.org/10.1007/s00221-005-2335-5

Cohen, Y., Schneidman, E., Paz, R., 2021. The geometry of neuronal representations during rule learning reveals complementary roles of cingulate cortex and putamen. Neuron 109, 839–851.e9. https://doi.org/10.1016/j.neuron.2020.12.027

Cromwell, H.C., Schultz, W., 2003. Effects of expectations for different reward magnitudes on neuronal activity in primate striatum. J. Neurophysiol. 89, 2823–2838. https://doi.org/10.1152/jn.01014.2002

Cummine, J., Cribben, I., Luu, C., Kim, E., Bakhtiari, R., Georgiou, G., Boliek, C.A., 2016. Understanding the Role of Speech Production in Reading: Evidence for a Print-to-Speech Neural Network Using Graphical Analysis. Neuropsychology. https://doi.org/10.1037/neu0000236

Cunnington, R., Windischberger, C., Deecke, L., Moser, E., 2002. The preparation and execution of self-initiated and externally-triggered movement: A study of event-related fMRI. Neuroimage 15, 373–385. https://doi.org/10.1006/nimg.2001.0976

Daw, N.D., Gershman, S.J., Seymour, B., Dayan, P., Dolan, R.J., 2011. Model-based influences on humans’ choices and striatal prediction errors. Neuron 69, 1204–1215. https://doi.org/10.1016/j.neuron.2011.02.027

Daw, N.D., Niv, Y., Dayan, P., 2005. Uncertainty-based competition between prefrontal and dorsolateral striatal systems for behavioral control. Nat. Neurosci. 8, 1704–1711. https://doi.org/10.1038/nn1560

De Houwer, J., 2019. On How Definitions of Habits Can Complicate Habit Research. Front. Psychol. 10, 1–9. https://doi.org/10.3389/fpsyg.2019.02642

de Wit, S., Barker, R. a, Dickinson, A.D., Cools, R., 2010. Habitual versus goal-directed action control in Parkinson disease. J Cogn Neurosci 23, 1218–1229. https://doi.org/10.1162/jocn.2010.21514 [doi]

de Wit, S., Kindt, M., Knot, S.L., Verhoeven, A.A.C., Robbins, T.W., Gasull-Camos, J., Evans, M., Mirza, H., Gillan, C.M., 2018. Shifting the balance between goals and habits: Five failures in experimental habit induction. J. Exp. Psychol. Gen. 147, 1043–1065. https://doi.org/10.1037/xge0000402

de Wit, S., Watson, P., Harsay, H.A., Cohen, M.X., van de Vijver, I., Ridderinkhof, K.R., 2012a. Corticostriatal connectivity underlies individual differences in the balance between habitual and goal-directed action control. J. Neurosci. 32, 12066–12075. https://doi.org/10.1523/JNEUROSCI.1088-12.2012

de Wit, S., Watson, P., Harsay, H.A., Cohen, M.X., van de Vijver, I., Ridderinkhof, K.R., 2012b. Corticostriatal connectivity underlies individual differences in the balance between habitual and goal-directed action control. J. Neurosci. 32, 12066–12075. https://doi.org/10.1523/JNEUROSCI.1088-12.2012

Delgado, M.R., Miller, M.M., Inati, S., Phelps, E.A., 2005. An fMRI study of reward-related probability learning. Neuroimage 24, 862–873. https://doi.org/10.1016/j.neuroimage.2004.10.002

Deserno, L., Huys, Q.J.M., Boehme, R., Buchert, R., Heinze, H.J., Grace, A.A., Dolan, R.J., Heinz, A., Schlagenhauf, F., 2015. Ventral striatal dopamine reflects behavioral and neural signatures of model-based control during sequential decision making. Proc. Natl. Acad. Sci. U. S. A. 112, 1595–1600. https://doi.org/10.1073/pnas.1417219112

Dickinson, A., Trans, P., Lond, R.S., 1985. Actions and habits: the development of behavioural autonomy. Philos. Trans. R. Soc. London. B, Biol. Sci. 308, 67–78. https://doi.org/10.1098/rstb.1985.0010

Doll, B.B., Duncan, K.D., Simon, D.A., Shohamy, D., Daw, N.D., 2015. Model-based choices involve prospective neural activity. Nat. Neurosci. 18, 767–772. https://doi.org/10.1038/NN.3981

Eickhoff, S.B., Laird, A.R., Grefkes, C., Wang, L.E., Zilles, K., Fox, P.T., 2009. Coordinate-based activation likelihood estimation meta-analysis of neuroimaging data: A random-effects approach based on empirical estimates of spatial uncertainty. Hum. Brain Mapp. 30, 2907–2926. https://doi.org/10.1002/hbm.20718

Eickhoff, S.B., Nichols, T.E., Laird, A.R., Hoffstaedter, F., Amunts, K., Fox, P.T., Bzdok, D., Eickhoff, C.R., 2016a. Behavior, sensitivity, and power of activation likelihood estimation characterized by massive empirical simulation. Neuroimage 137, 70–85. https://doi.org/10.1016/j.neuroimage.2016.04.072

Eickhoff, S.B., Nichols, T.E., Laird, A.R., Hoffstaedter, F., Amunts, K., Fox, P.T., Bzdok, D., Eickhoff, C.R., 2016b. Behavior, sensitivity, and power of activation likelihood estimation characterized by massive empirical simulation. Neuroimage 137, 70–85. https://doi.org/10.1016/J.NEUROIMAGE.2016.04.072

Erdeniz, B., Done, J., 2019. Common and distinct functional brain networks for intuitive and deliberate decision making. Brain Sci. 9. https://doi.org/10.3390/brainsci9070174

Erhard, K., Kessler, F., Neumann, N., Ortheil, H.J., Lotze, M., 2014. Professional training in creative writing is associated with enhanced fronto-striatal activity in a literary text continuation task. Neuroimage 100, 15–23. https://doi.org/10.1016/j.neuroimage.2014.05.076

Eryilmaz, H., Rodriguez-Thompson, A., Tanner, A.S., Giegold, M., Huntington, F.C., Roffman, J.L., 2017. Neural determinants of human goal-directed vs. habitual action control and their relation to trait motivation. Sci. Rep. 7, 1–11. https://doi.org/10.1038/s41598-017-06284-y

Etchamendy, N., Konishi, K., Pike, G.B., Marighetto, A., Bohbot, V.D., 2012. Evidence for a virtual human analog of a rodent relational memory task: A study of aging and fMRI in young adults. Hippocampus 22, 869–880. https://doi.org/10.1002/HIPO.20948

Everitt, B.J., Belin, D., Economidou, D., Pelloux, Y., Dalley, J.W., Robbins, T.W., 2008. Neural mechanisms underlying the vulnerability to develop compulsive drug-seeking habits and addiction. Philos. Trans. R. Soc. B Biol. Sci. 363, 3125–3135. https://doi.org/10.1098/rstb.2008.0089

Fera, F., Weickert, T.W., Goldberg, T.E., Tessitore, A., Hariri, A., Das, S., Lee, S., Zoltick, B., Meeter, M., Myers, C.E., Gluck, M.A., Weinberger, D.R., Mattay, V.S., 2005. Neural mechanisms underlying probabilistic category learning in normal aging. J. Neurosci. 25, 11340–11348. https://doi.org/10.1523/JNEUROSCI.2736-05.2005

Fermin, A.S.R., Yoshida, T., Yoshimoto, J., Ito, M., Tanaka, S.C., Doya, K., 2016. Model-based action planning involves cortico-cerebellar and basal ganglia networks. Sci. Rep. 6. https://doi.org/10.1038/SREP31378

Fernández-Seara, M.A., Aznárez-Sanado, M., Mengual, E., Loayza, F.R., Pastor, M.A., 2009. Continuous performance of a novel motor sequence leads to highly correlated striatal and hippocampal perfusion increases. Neuroimage 47, 1797–1808. https://doi.org/10.1016/J.NEUROIMAGE.2009.05.061

Foerde, K., 2018. What are habits and do they depend on the striatum? A view from the study of neuropsychological populations. Curr. Opin. Behav. Sci. 20, 17–24. https://doi.org/10.1016/j.cobeha.2017.08.011

Foerde, K., Knowlton, B.J., Poldrack, R.A., 2006. Modulation of competing memory systems by distraction. Proc. Natl. Acad. Sci. U. S. A. 103, 11778–11783. https://doi.org/10.1073/PNAS.0602659103

Forstmann, B.U., Dutilh, G., Brown, S., Neumann, J., Von Cramon, D.Y., Ridderinkhof, K.R., Wagenmakers, E.J., 2008. Striatum and pre-SMA facilitate decision-making under time pressure. Proc. Natl. Acad. Sci. U. S. A. 105, 17538–17542. https://doi.org/10.1073/pnas.0805903105

Fox, P.T., Laird, A.R., Eickhoff, S.B., Lancaster, J.L., Fox, M., Uecker, A.M., Robertson, M., Ray, K.L., 2013. User manual for GingerALE 2.3, Retrieved from http://brainmap.org/ale/manual.pdf.

Gardiner, T.W., Nelson, R.J., 1992. Striatal neuronal activity during the initiation and execution of hand movements made in response to visual and vibratory cues. Exp. Brain Res. 92, 15–26. https://doi.org/10.1007/BF00230379

Gardner, B., 2015. A review and analysis of the use of “habit” in understanding, predicting and influencing health-related behaviour. Health Psychol. Rev. 9, 277–295. https://doi.org/10.1080/17437199.2013.876238

Gillan, C.M., 2021. Recent Developments in the Habit Hypothesis of OCD and Compulsive Disorders. Springer, Berlin, Heidelberg, pp. 1–21. https://doi.org/10.1007/7854_2020_199

Gläscher, J., Daw, N., Dayan, P., O’Doherty, J.P., 2010. States versus rewards: Dissociable neural prediction error signals underlying model-based and model-free reinforcement learning. Neuron 66, 585–595. https://doi.org/10.1016/j.neuron.2010.04.016

Gottfried, J.A., O’Doherty, J., Dolan, R.J., 2003. Encoding predictive reward value in human amygdala and orbitofrontal cortex. Science (80-.). 301, 1104–1107. https://doi.org/10.1126/SCIENCE.1087919

Gould, L., Mickleborough, M.J.S., Lorentz, E., Ekstrand, C., Borowsky, R., 2018. A behavioral and fMRI examination of the effect of rhythm on reading noun-verb homographs aloud. https://doi.org/10.1080/23273798.2018.1442012 33, 829–849. https://doi.org/10.1080/23273798.2018.1442012

Grabenhorst, F., Rolls, E.T., 2011. Value, pleasure and choice in the ventral prefrontal cortex. Trends Cogn. Sci. 15, 56–67. https://doi.org/10.1016/j.tics.2010.12.004

Graydon, F.X., Young, R., Benton, M.D., Genik, R.J., Posse, S., Hsieh, L., Green, C., 2004. Visual event detection during simulated driving: Identifying the neural correlates with functional neuroimaging. Transp. Res. Part F Traffic Psychol. Behav. 7, 271–286. https://doi.org/10.1016/j.trf.2004.09.006

Haber, S.N., Fudge, J.L., McFarland, N.R., 2000. Striatonigrostriatal Pathways in Primates Form an Ascending Spiral from the Shell to the Dorsolateral Striatum. J. Neurosci. 20, 2369–2382. https://doi.org/10.1523/JNEUROSCI.20-06-02369.2000

Haber, S.N., Knutson, B., 2009. The Reward Circuit: Linking Primate Anatomy and Human Imaging. Neuropsychopharmacology 35, 4–26. https://doi.org/10.1038/npp.2009.129

Haith, A.M., Krakauer, J.W., 2018. The multiple effects of practice: skill, habit and reduced cognitive load. Curr. Opin. Behav. Sci. 20, 196–201. https://doi.org/10.1016/j.cobeha.2018.01.015

Hardwick, R.M., Forrence, A.D., Krakauer, J.W., Haith, A.M., 2019. Time-dependent competition between goal-directed and habitual response preparation. Nat. Hum. Behav. 3, 1252–1262. https://doi.org/10.1038/s41562-019-0725-0

Haruno, M., Kawato, M., 2006. Different neural correlates of reward expectation and reward expectation error in the putamen and caudate nucleus during stimulus-action-reward association learning. J. Neurophysiol. 95, 948–959. https://doi.org/10.1152/jn.00382.2005

Haynes, W.I.A., Haber, S.N., 2013. The organization of prefrontal-subthalamic inputs in primates provides an anatomical substrate for both functional specificity and integration: Implications for basal ganglia models and deep brain stimulation. J. Neurosci. 33, 4804–4814. https://doi.org/10.1523/JNEUROSCI.4674-12.2013

Hernandez, L.F., Obeso, I., Costa, R.M., Redgrave, P., Obeso, J.A., 2019. Dopaminergic Vulnerability in Parkinson Disease: The Cost of Humans’ Habitual Performance. Trends Neurosci. 42, 375–383. https://doi.org/10.1016/j.tins.2019.03.007

Hikosaka, O., Takikawa, Y., Kawagoe, R., Kojima, S., Doupe, A.J., Silver, R., Boahen, K., Grillner, S., Kopell, N., Olsen, K.L., Neurosci F-W Zhou, J., Matta, S.G., Zhou, F., Neurophysiol, J., Bargas Carrillo-Reid, J.L., Tecuapetla, F., Tapia, D., Hernandez-Cruz, A., Galarraga, E., Drucker-Colin, R., 2000. Tutor Song Exposure Song Selectivity in the Pallial-Basal Ganglia Song Circuit of Zebra Finches Raised Without Role of the Basal Ganglia in the Control of Purposive Saccadic Eye Movements. Physiol Rev 80, 953– 978.

Holl, A.K., Wilkinson, L., Tabrizi, S.J., Painold, A., Jahanshahi, M., 2012. Probabilistic classification learning with corrective feedback is selectively impaired in early Huntington’s disease--evidence for the role of the striatum in learning with feedback. Neuropsychologia 50, 2176–2186. https://doi.org/10.1016/J.NEUROPSYCHOLOGIA.2012.05.021

Horovitz, S.G., Gallea, C., Najee-ullah, M.A., Hallett, M., 2013. Functional Anatomy of Writing with the Dominant Hand. PLoS One 8, 1–10. https://doi.org/10.1371/journal.pone.0067931

Hsieh, L., Young, R.A., Bowyer, S.M., Moran, J.E., Genik, R.J., Green, C.C., Chiang, Y.R., Yu, Y.J., Liao, C.C., Seaman, S., 2009. Conversation effects on neural mechanisms underlying reaction time to visual events while viewing a driving scene: fMRI analysis and asynchrony model. Brain Res. https://doi.org/10.1016/j.brainres.2008.10.002

Hsu, C.T., Jacobs, A.M., Citron, F.M.M., Conrad, M., 2015. The emotion potential of words and passages in reading Harry Potter - An fMRI study. Brain Lang. 142, 96–114. https://doi.org/10.1016/j.bandl.2015.01.011

Hu, X.Y., Wang, L., Liu, H., Zhang, S.Z., 2006. Functional magnetic resonance imaging study of writer’s cramp. Chin. Med. J. (Engl). 119, 1263–1271. https://doi.org/10.1097/00029330-200608010-00006

Huang, Y., Yaple, Z.A., Yu, R., 2020. Goal-oriented and habitual decisions: Neural signatures of model-based and model-free learning. Neuroimage 215, 116834. https://doi.org/10.1016/j.neuroimage.2020.116834

Huth, A.G., De Heer, W.A., Griffiths, T.L., Theunissen, F.E., Gallant, J.L., 2016. Natural speech reveals the semantic maps that tile human cerebral cortex. Nature 532, 453–458. https://doi.org/10.1038/nature17637

Iaria, G., Petrides, M., Dagher, A., Pike, B., Bohbot, V.D., 2003. Cognitive strategies dependent on the hippocampus and caudate nucleus in human navigation: variability and change with practice. J. Neurosci. Off. J. Soc. Neurosci. 23, 5945–5952. https://doi.org/10.1523/jneurosci.23-13-05945.2003

Jaeger L, Marchal-Crespo L, Wolf P, Riener R, Michels L, K.S., 2014. Brain activation associated with active and passive lower limb stepping. Front. Hum. Neurosci. 8. https://doi.org/10.3389/fnhum.2014.00828

Jankowski, J., Scheef, L., Hüppe, C., Boecker, H., 2009. Distinct striatal regions for planning and executing novel and automated movement sequences. Neuroimage 44, 1369– 1379. https://doi.org/10.1016/j.neuroimage.2008.10.059

Jeanvoine, H., Labriffe, M., Tannou, T., Navasiolava, N., Ter Minassian, A., Girot, J.B., Leiber, L.M., Custaud, M.A., Annweiler, C., Dinomais, M., 2022. Specific age-correlated activation of top hierarchical motor control areas during gait-like plantar stimulation: An fMRI study. Hum. Brain Mapp. 43, 833–843. https://doi.org/10.1002/HBM.25691

Karimpoor, M., Churchill, N.W., Tam, F., Fischer, C.E., Schweizer, T.A., Graham, S.J., 2018. Functional MRI of handwriting tasks: A study of healthy young adults interacting with a novel touch-sensitive tablet. Front. Hum. Neurosci. 12, 1–14. https://doi.org/10.3389/fnhum.2018.00030

Karimpoor, M., Tam, F., Strother, S.C., Fischer, C.E., Schweizer, T.A., Graham, S.J., 2015. A computerized tablet with visual feedback of hand position for functional magnetic resonance imaging. Front. Hum. Neurosci. 9, 150. https://doi.org/10.3389/fnhum.2015.00150

Katanoda, K., Yoshikawa, K., Sugishita, M., 2001. A functional MRI study on the neural substrates for writing. Hum. Brain Mapp. 13, 34–42. https://doi.org/10.1002/hbm.1023

Kilner, J.M., Neal, A., Weiskopf, N., Friston, K.J., Frith, C.D., 2009. Evidence of Mirror Neurons in Human Inferior Frontal Gyrus. J. Neurosci. 29, 10153. https://doi.org/10.1523/JNEUROSCI.2668-09.2009

Kim, W., Chang, Y., Kim, J., Seo, J., Ryu, K., Lee, E., Woo, M., Janelle, C.M., 2014. An fMRI study of differences in brain activity among elite, expert, and novice archers at the moment of optimal aiming. Cogn. Behav. Neurol. 27, 173–182. https://doi.org/10.1097/WNN.0000000000000042

Kimura, M., 1986. The role of primate putamen neurons in the association of sensory stimuli with movement. Neurosci. Res. 3, 436–443. https://doi.org/10.1016/0168-0102(86)90035-0

Kimura, M., Kato, M., Shimazaki, H., 1990. Physiological properties of projection neurons in the monkey striatum to the globus pallidus. Exp. Brain Res. 82, 672–676. https://doi.org/10.1007/BF00228811

Kimura, M., Rajkowski, J., Evarts, E., 1984. Tonically discharging putamen neurons exhibit set-dependent responses. Proc. Natl. Acad. Sci. U. S. A. 81, 4998–5001. https://doi.org/10.1073/pnas.81.15.4998

Kunimatsu, J., Maeda, K., Hikosaka, O., 2019. The caudal part of putamen represents the historical object value information. J. Neurosci. 39, 1709–1719. https://doi.org/10.1523/JNEUROSCI.2534-18.2018

la Fougère, C., Zwergal, A., Rominger, A., Förster, S., Fesl, G., Dieterich, M., Brandt, T., Strupp, M., Bartenstein, P., Jahn, K., 2010. Real versus imagined locomotion: A [18F]-FDG PET-fMRI comparison. Neuroimage 50, 1589–1598. https://doi.org/10.1016/j.neuroimage.2009.12.060

Lancaster, J.L., Tordesillas-Gutiérrez, D., Martinez, M., Salinas, F., Evans, A., Zilles, K., Mazziotta, J.C., Fox, P.T., 2007. Bias between MNI and talairach coordinates analyzed using the ICBM-152 brain template. Hum. Brain Mapp. 28, 1194–1205. https://doi.org/10.1002/hbm.20345

Leh, S.E., Ptito, A., Chakravarty, M.M., Strafella, A.P., 2007. Fronto-striatal connections in the human brain: A probabilistic diffusion tractography study. Neurosci. Lett. 419, 113– 118. https://doi.org/10.1016/j.neulet.2007.04.049

Lehéricy, S., Benali, H., Van De Moortele, P.F., Pélégrini-Issac, M., Waechter, T., Ugurbil, K., Doyon, J., 2005a. Distinct basal ganglia territories are engaged in early and advanced motor sequence learning. Proc. Natl. Acad. Sci. U. S. A. 102, 12566–12571. https://doi.org/10.1073/pnas.0502762102

Lehéricy, S., Benali, H., Van De Moortele, P.F., Pélégrini-Issac, M., Waechter, T., Ugurbil, K., Doyon, J., 2005b. Distinct basal ganglia territories are engaged in early and advanced motor sequence learning. Proc. Natl. Acad. Sci. U. S. A. 102, 12566–12571. https://doi.org/10.1073/pnas.0502762102

Leisman, G., Moustafa, A., Shafir, T., 2016. Thinking, Walking, Talking: Integratory Motor and Cognitive Brain Function. Front. Public Heal. 4, 1. https://doi.org/10.3389/fpubh.2016.00094

Liljeholm, M., Dunne, S., O’Doherty, J.P., 2015a. Differentiating neural systems mediating the acquisition vs. expression of goal-directed and habitual behavioral control. Eur. J. Neurosci. 41, 1358–1371. https://doi.org/10.1111/ejn.12897

Liljeholm, M., Dunne, S., O’Doherty, J.P., 2015b. Differentiating neural systems mediating the acquisition vs. expression of goal-directed and habitual behavioral control. Eur. J. Neurosci. 41, 1358–1371. https://doi.org/10.1111/ejn.12897

Liljeholm, M., O’doherty, J.P., 2012. Contributions of the striatum to learning, motivation, and performance: an associative account. Trends Cogn Sci 16, 467–475. https://doi.org/10.1016/j.tics.2012.07.007

Lin, Z., Tam, F., Churchill, N.W., Schweizer, T.A., Graham, S.J., 2021. Tablet technology for writing and drawing during functional magnetic resonance imaging: A review. Sensors (Switzerland). https://doi.org/10.3390/s21020401

Longcamp, M., Lagarrigue, A., Nazarian, B., Roth, M., Anton, J.L., Alario, F.X., Velay, J.L., 2014. Functional specificity in the motor system: Evidence from coupled fMRI and kinematic recordings during letter and digit writing. Hum. Brain Mapp. 35, 6077–6087. https://doi.org/10.1002/hbm.22606

Luque, D., Beesley, T., Morris, R.W., Jack, B.N., Griffiths, O., Whitford, T.J., Le Pelley, M.E., 2017. Goal-Directed and Habit-Like Modulations of Stimulus Processing during Reinforcement Learning. https://doi.org/10.1523/JNEUROSCI.3205-16.2017

Luque, D., Molinero, S., Watson, P., López, F.J., Le Pelley, M.E., 2020. Measuring habit formation through goal-directed response switching. J. Exp. Psychol. Gen. 149, 1449– 1459. https://doi.org/10.1037/xge0000722

Marchal, V., Sellers, J., Pélégrini-Issac, M., Galléa, C., Bertasi, E., Valabrègue, R., Lau, B., Leboucher, P., Bardinet, E., Welter, M.L., Karachi, C., 2019. Deep brain activation patterns involved in virtual gait without and with a doorway: An fMRI study. PLoS One 14. https://doi.org/10.1371/journal.pone.0223494

Martínez, M., Valencia, M., Vidorreta, M., Luis, E.O., Castellanos, G., Villagra, F., Fern?? ndez-Seara, M.A., Pastor, M.A., 2016a. Trade-off between frequency and precision during stepping movements: Kinematic and BOLD brain activation patterns. Hum. Brain Mapp. 37, 1722–1737. https://doi.org/10.1002/hbm.23131

Martínez, M., Valencia, M., Vidorreta, M., Luis, E.O., Castellanos, G., Villagra, F., Fernández-Seara, M.A., Pastor, M.A., 2016b. Trade-off between frequency and precision during stepping movements: Kinematic and BOLD brain activation patterns. Hum. Brain Mapp. 37, 1722–1737. https://doi.org/10.1002/hbm.23131

Martínez, M., Villagra, F., Castellote, J.M., Pastor, M.A., 2018. Kinematic and kinetic patterns related to free-walking in parkinson’s disease. Sensors (Switzerland) 18. https://doi.org/10.3390/s18124224

Martinez, M., Villagra, F., Loayza, F., Vidorreta, M., Arrondo, G., Luis, E., Diaz, J., Echeverria, M., Fernandez-Seara, M.A., Pastor, M.A., 2014. MRI-compatible device for examining brain activation related to stepping. IEEE Trans. Med. Imaging 33, 1044– 1053. https://doi.org/10.1109/TMI.2014.2301493

Mas-Herrero, E., Sescousse, G., Cools, R., Marco-Pallarés, J., 2019. The contribution of striatal pseudo-reward prediction errors to value-based decision-making. Neuroimage 193, 67–74. https://doi.org/10.1016/j.neuroimage.2019.02.052

Matsuda, W., Furuta, T., Nakamura, K.C., Hioki, H., Fujiyama, F., Arai, R., Kaneko, T., 2009. Single nigrostriatal dopaminergic neurons form widely spread and highly dense axonal arborizations in the neostriatum. J. Neurosci. 29, 444–453. https://doi.org/10.1523/JNEUROSCI.4029-08.2009

Mazziotta, J., Toga, A., Evans, A., Fox, P., Lancaster, J., Zilles, K., Woods, R., Paus, T., Simpson, G., Pike, B., Holmes, C., Collins, L., Thompson, P., MacDonald, D., Iacoboni, M., Schormann, T., Amunts, K., Palomero-Gallagher, N., Geyer, S., Parsons, L., Narr, K., Kabani, N., Le Goualher, G., Boomsma, D., Cannon, T., Kawashima, R., Mazoyer, B., 2001. A probabilistic atlas and reference system for the human brain: International Consortium for Brain Mapping (ICBM). Philos. Trans. R. Soc. B Biol. Sci. https://doi.org/10.1098/rstb.2001.0915

McFarland, N.R., Haber, S.N., 2002. Thalamic relay nuclei of the basal ganglia form both reciprocal and nonreciprocal cortical connections, linking multiple frontal cortical areas. J. Neurosci. 22, 8117–8132. https://doi.org/10.1523/jneurosci.22-18-08117.2002

Mechelli, A., Gorno-Tempini, M.L., Price, C.J., 2003. Neuroimaging studies of word and pseudoword reading: Consistencies, inconsistencies, and limitations. J. Cogn. Neurosci. 15, 260–271. https://doi.org/10.1162/089892903321208196

Meschyan, G., Hernandez, A.E., 2006. Impact of language proficiency and orthographic transparency on bilingual word reading: An fMRI investigation. Neuroimage 29, 1135– 1140. https://doi.org/10.1016/j.neuroimage.2005.08.055

Mi, T.M., Zhang, W., McKeown, M.J., Chan, P., 2021. Impaired Formation and Expression of Goal-Directed and Habitual Control in Parkinson’s Disease. Front. Aging Neurosci. 13. https://doi.org/10.3389/FNAGI.2021.734807

Miller, K.J., Shenhav, A., Ludvig, E.A., 2019. Habits without values. Psychol. Rev. https://doi.org/10.1037/rev0000120

Molenberghs, P., Cunnington, R., Mattingley, J.B., 2012. Brain regions with mirror properties: A meta-analysis of 125 human fMRI studies. Neurosci. Biobehav. Rev. 36, 341–349. https://doi.org/10.1016/J.NEUBIOREV.2011.07.004

Moore, C.J., Price, C.J., 1999. Three distinct ventral occipitotemporal regions for reading and object naming. Neuroimage 10, 181–192. https://doi.org/10.1006/nimg.1999.0450

Morris, L.S., Kundu, P., Dowell, N., Mechelmans, D.J., Favre, P., Irvine, M.A., Robbins, T.W., Daw, N., Bullmore, E.T., Harrison, N.A., Voon, V., 2016. Fronto-striatal organization: Defining functional and microstructural substrates of behavioural flexibility. Cortex 74, 118–133. https://doi.org/10.1016/j.cortex.2015.11.004

Morris, N.J., Stein, C.M., 2017. Model-free linkage analysis of a quantitative trait, in: Methods in Molecular Biology. Humana Press Inc., pp. 327–342. https://doi.org/10.1007/978-1-4939-7274-6_16

Müller, V.I., Cieslik, E.C., Laird, A.R., Fox, P.T., Radua, J., Mataix-Cols, D., Tench, C.R., Yarkoni, T., Nichols, T.E., Turkeltaub, P.E., Wager, T.D., Eickhoff, S.B., 2018. Ten simple rules for neuroimaging meta-analysis. Neurosci. Biobehav. Rev. 84, 151. https://doi.org/10.1016/J.NEUBIOREV.2017.11.012

Murphy, K., Garavan, H., 2004. An empirical investigation into the number of subjects required for an event-related fMRI study. Neuroimage 22, 879–885. https://doi.org/10.1016/J.NEUROIMAGE.2004.02.005

Nachev, P., Kennard, C., Husain, M., 2008. Functional role of the supplementary and pre-supplementary motor areas. Nat. Rev. Neurosci. https://doi.org/10.1038/nrn2478

Nakamura, K., Honda, M., Hirano, S., Oga, T., Sawamoto, N., Hanakawa, T., Inoue, H., Ito, J., Matsuda, T., Fukuyama, H., Shibasaki, H., 2002. Modulation of the visual word retrieval system in writing: A functional MRI study on the Japanese orthographies. J. Cogn. Neurosci. 14, 104–115. https://doi.org/10.1162/089892902317205366

Nebe, S., Kroemer, N.B., Schad, D.J., Bernhardt, N., Sebold, M., Müller, D.K., Scholl, L., Kuitunen-Paul, S., Heinz, A., Rapp, M.A., Huys, Q.J.M., Smolka, M.N., 2018. No association of goal-directed and habitual control with alcohol consumption in young adults. Addict. Biol. 23, 379–393. https://doi.org/10.1111/ADB.12490

Noble, J.W., Eng, J.J., Boyd, L.A., 2014. Bilateral motor tasks involve more brain regions and higher neural activation than unilateral tasks: An fMRI study. Exp. Brain Res. 232, 2785–2795. https://doi.org/10.1007/s00221-014-3963-4

Oberhuber, M., Hope, T.M.H., Seghier, M.L., Parker Jones, O., Prejawa, S., Green, D.W., Price, C.J., 2016. Four Functionally Distinct Regions in the Left Supramarginal Gyrus Support Word Processing. Cereb. Cortex 26, 4212–4226. https://doi.org/10.1093/cercor/bhw251

Oberhuber, M., Jones, O.P., Hope, T.M.H., Prejawa, S., Seghier, M.L., Green, D.W., Price, C.J., 2013. Functionally distinct contributions of the anterior and posterior putamen during sublexical and lexical reading. Front. Hum. Neurosci. https://doi.org/10.3389/fnhum.2013.00787

Oh-Descher, H., Beck, J.M., Ferrari, S., Sommer, M.A., Egner, T., 2017. Probabilistic inference under time pressure leads to a cortical-to-subcortical shift in decision evidence integration. Neuroimage 162, 138–150. https://doi.org/10.1016/j.neuroimage.2017.08.069

Ohbayashi, M., Picard, N., Strick, P.L., 2016. Inactivation of the dorsal premotor area disrupts internally generated, but not visually guided, sequential movements. J. Neurosci. 36, 1971–1976. https://doi.org/10.1523/JNEUROSCI.2356-15.2016

Parkes, S.L., Bradfield, L.A., Balleine, B.W., 2015. Interaction of insular cortex and ventral striatum mediates the effect of incentive memory on choice between goal-directed actions. J. Neurosci. 35, 6464–6471. https://doi.org/10.1523/JNEUROSCI.4153-14.2015

Patterson, T.K., Knowlton, B.J., 2018. Subregional specificity in human striatal habit learning: a meta-analytic review of the fMRI literature. Curr. Opin. Behav. Sci. 20, 75–82. https://doi.org/10.1016/j.cobeha.2017.10.005

Perez, Omar D, Dickinson, A., 2020. A theory of actions and habits: The interaction of rate correlation and contiguity systems in free-operant behavior. https://doi.org/10.1037/rev0000201

Perez, Omar D., Dickinson, A., 2020. A theory of actions and habits: The interaction of rate correlation and contiguity systems in free-operant behavior. Psychol. Rev. 127, 945– 971. https://doi.org/10.1037/REV0000201

Peters, A.J., Fabre, J.M.J., Steinmetz, N.A., Harris, K.D., Carandini, M., 2021. Striatal activity topographically reflects cortical activity. Nature 591, 420–425. https://doi.org/10.1038/s41586-020-03166-8

Peters, S., Eng, J.J., Liu-Ambrose, T., Borich, M.R., Dao, E., Amanian, A., Boyd, L.A., 2019. Brain activity associated with Dual-task performance of Ankle motor control during cognitive challenge. Brain Behav. 9. https://doi.org/10.1002/brb3.1349

Peterson, B.S., Riddle, M.A., Cohen, D.J., Katz, L.D., Smith, J.C., Leckman, J.F., 1993. Human basal ganglia volume asymmetries on magnetic resonance images. Magn. Reson. Imaging 11, 493–498. https://doi.org/10.1016/0730-725X(93)90468-S

Peterson, D.S., Horak, F.B., 2016. Neural control of walking in people with parkinsonism. Physiology. https://doi.org/10.1152/physiol.00034.2015

Pineda-Pardo, J.D.S.A., Sánchez-Ferro, Á., Monje, M.H.G., Pavese, N., Obeso, J.A., 2022. Onset pattern of nigrostriatal denervation in early Parkinson’s disease. Brain 145, 1018–1028. https://doi.org/10.1093/BRAIN/AWAB378

Poldrack, R.A., Clark, J., Paré-Blagoev, E.J., Shohamy, D., Creso Moyano, J., Myers, C., Gluck, M.A., 2001. Interactive memory systems in the human brain. Nature 414, 546–550. https://doi.org/10.1038/35107080

Poldrack, R.A., Prabhakaran, V., Seger, C.A., Gabrieli, J.D.E., 1999. Striatal activation during acquisition of a cognitive skill. Neuropsychology 13, 564–574. https://doi.org/10.1037//0894-4105.13.4.564

Pool, E., Gera, R., Fransen, A., Perez, O.D., Cremer, A., Aleksic, M., Tanwisuth, S., Quail, S., Ceceli, A.O., Manfredi, D., Nave, G., Tricomi, E., Balleine, B., Schonberg, T., Schwabe, L., O’Doherty, J.P., n.d. Determining the effects of training duration on the behavioral expression of habitual control in humans: a multi-laboratory investigation. https://doi.org/10.31234/OSF.IO/Z756H

Potgieser, A.R.E., Van Der Hoorn, A., De Jong, B.M., 2015. Cerebral Activations Related to Writing and Drawing with each hand. PLoS One 10, 1–25. https://doi.org/10.1371/journal.pone.0126723

Redgrave, P., Gurney, K., Reynolds, J., 2008. What is reinforced by phasic dopamine signals? Brain Res. Rev. https://doi.org/10.1016/j.brainresrev.2007.10.007

Redgrave, P., Prescott, T.J., Gurney, K., 1999. The basal ganglia: a vertebrate solution to the selection problem? Neuroscience 89, 1009–1023. https://doi.org/S0306452298003194 [pii]

Redgrave, P., Rodriguez, M., Smith, Y., Rodriguez-Oroz, M.C., Lehericy, S., Bergman, H., Agid, Y., Delong, M.R., Obeso, J.A., 2010a. Goal-directed and habitual control in the basal ganglia: Implications for Parkinson’s disease. Nat. Rev. Neurosci. 11, 760–772. https://doi.org/10.1038/nrn2915

Redgrave, P., Rodriguez, M., Smith, Y., Rodriguez-Oroz, M.C., Lehericy, S., Bergman, H., Agid, Y., Delong, M.R., Obeso, J.A., 2010b. Goal-directed and habitual coRedgrave, P., Rodriguez, M., Smith, Y., Rodriguez-Oroz, M. C., Lehericy, S., Bergman, H., Agid, Y., Delong, M. R., & Obeso, J. A. (2010). Goal-directed and habitual control in the basal ganglia: Implications for Parkinson’s di. Nat. Rev. Neurosci. 11, 760–772. https://doi.org/10.1038/nrn2915

Romero, M.C., Bermudez, M.A., Vicente, A.F., Perez, R., Gonzalez, F., 2008. Activity of neurons in the caudate and putamen during a visuomotor task. Neuroreport 19, 1141– 1145. https://doi.org/10.1097/WNR.0b013e328307c3fc

Rueckl, J.G., Paz-Alonso, P.M., Molfese, P.J., Kuo, W.J., Bick, A., Frost, S.J., Hancock, R., Wu, D.H., Einar Mencl, W., Duñabeitia, J.A., Lee, J.R., Oliver, M., Zevin, J.D., Hoeft, F., Carreiras, M., Tzeng, O.J.L., Pugh, K.R., Frost, R., 2015. Universal brain signature of proficient reading: Evidence from four contrasting languages. Proc. Natl. Acad. Sci. U. S. A. 112, 15510–15515. https://doi.org/10.1073/pnas.1509321112

Sauvage, C., Jissendi, P., Seignan, S., Manto, M., Habas, C., 2013. Brain areas involved in the control of speed during a motor sequence of the foot: Real movement versus mental imagery. J. Neuroradiol. 40, 267–280. https://doi.org/10.1016/j.neurad.2012.10.001

Schultz, W., 2006. Behavioral theories and the neurophysiology of reward. Annu. Rev. Psychol. 57, 87–115. https://doi.org/10.1146/annurev.psych.56.091103.070229

Schwabe, L., Tegenthoff, M., Höffken, O., Wolf, O.T., 2013. Mineralocorticoid Receptor Blockade Prevents Stress-Induced Modulation of Multiple Memory Systems in the Human Brain. Biol. Psychiatry 74, 801–808. https://doi.org/10.1016/J.BIOPSYCH.2013.06.001

Segal, E., Petrides, M., 2012. The anterior superior parietal lobule and its interactions with language and motor areas during writing. Eur. J. Neurosci. 35, 309–322. https://doi.org/10.1111/j.1460-9568.2011.07937.x

Seghier, M.L., Lee, H.L., Schofield, T., Ellis, C.L., Price, C.J., 2008. Inter-subject variability in the use of two different neuronal networks for reading aloud familiar words. Neuroimage 42, 1226–1236. https://doi.org/10.1016/j.neuroimage.2008.05.029

Seghier, M.L., Price, C.J., 2010. Reading Aloud Boosts Connectivity through the Putamen. Cereb. Cortex 20, 570–582. https://doi.org/10.1093/cercor/bhp123

Sescousse, G., Caldú, X., Segura, B., Dreher, J.C., 2013. Processing of primary and secondary rewards: A quantitative meta-analysis and review of human functional neuroimaging studies. Neurosci. Biobehav. Rev. 37, 681–696. https://doi.org/10.1016/j.neubiorev.2013.02.002

Sjoerds, Z., De Wit, S., Van Den Brink, W., Robbins, T.W., Beekman, A.T.F., Penninx, B.W.J.H., Veltman, D.J., 2013. Behavioral and neuroimaging evidence for overreliance on habit learning in alcohol-dependent patients. Transl. Psychiatry 3, e337–8. https://doi.org/10.1038/tp.2013.107

Smith, K.S., Graybiel, A.M., 2016. Habit formation. Dialogues Clin. Neurosci. 18, 33–43. https://doi.org/10.31887/dcns.2016.18.1/ksmith

Smittenaar, P., Guitart-Masip, M., Lutti, A., Dolan, R.J., 2013. Preparing for selective inhibition within frontostriatal loops. J. Neurosci. 33, 18087–18097. https://doi.org/10.1523/JNEUROSCI.2167-13.2013

Soares, J.M., Sampaio, A., Ferreira, L.M., Santos, N.C., Marques, F., Palha, J.A., Cerqueira, J.J., Sousa, N., 2012. Stress-induced changes in human decision-making are reversible. Transl. Psychiatry 2. https://doi.org/10.1038/TP.2012.59

Sommer, S., Pollmann, S., 2016. Putamen Activation Represents an Intrinsic Positive Prediction Error Signal for Visual Search in Repeated Configurations. Open Neuroimag. J. 10, 126–138. https://doi.org/10.2174/1874440001610010126

Spiers, H.J., Maguire, E.A., 2007. Neural substrates of driving behaviour. Neuroimage 36, 245–255. https://doi.org/10.1016/j.neuroimage.2007.02.032

Steele, C.J., Penhune, V.B., 2010. Specific Increases within Global Decreases: A Functional Magnetic Resonance Imaging Investigation of Five Days of Motor Sequence Learning. J. Neurosci. 30, 8332–8341. https://doi.org/10.1523/JNEUROSCI.5569-09.2010

Sun, R., 2004. Desiderata for cognitive architectures. Philos. Psychol. 17, 341–373. https://doi.org/10.1080/0951508042000286721

Swett, B.A., Contreras-Vidal, J.L., Birn, R., Braun, A., 2010. Neural Substrates of Graphomotor Sequence Learning: A Combined fMRI and Kinematic Study. J. Neurophysiol. 103, 3366. https://doi.org/10.1152/JN.00449.2009

Swinnen, S.P., Vangheluwe, S., Wagemans, J., Coxon, J.P., Goble, D.J., Van Impe, A., Sunaert, S., Peeters, R., Wenderoth, N., 2010. Shared neural resources between left and right interlimb coordination skills: The neural substrate of abstract motor representations. Neuroimage 49, 2570–2580. https://doi.org/10.1016/j.neuroimage.2009.10.052

Tanaka, S.C., Balleine, B.W., O’Doherty, J.P., 2008. Calculating consequences: Brain systems that encode the causal effects of actions. J. Neurosci. 28, 6750–6755. https://doi.org/10.1523/JNEUROSCI.1808-08.2008

Toyomura, A., Shibata, M., Kuriki, S., 2012. Self-paced and externally triggered rhythmical lower limb movements: A functional MRI study. Neurosci. Lett. 516, 39–44. https://doi.org/10.1016/j.neulet.2012.03.049

Tricomi, E., Balleine, B.W., O’Doherty, J.P., 2009. A specific role for posterior dorsolateral striatum in human habit learning. Eur. J. Neurosci. 29, 2225–2232. https://doi.org/10.1111/j.1460-9568.2009.06796.x

Trinastic, J.P., Kautz, S.A., McGregor, K., Gregory, C., Bowden, M., Benjamin, M.B., Kurtzman, M., Chang, Y.L., Conway, T., Crosson, B., 2010. An fMRI study of the differences in brain activity during active ankle dorsiflexion and plantarflexion. Brain Imaging Behav. 4, 121–131. https://doi.org/10.1007/s11682-010-9091-2

Tunik, E., Houk, J.C., Grafton, S.T., 2009. Basal ganglia contribution to the initiation of corrective submovements. Neuroimage 47, 1757–1766. https://doi.org/10.1016/j.neuroimage.2009.04.077

Turkeltaub, P.E., Eden, G.F., Jones, K.M., Zeffiro, T.A., 2002. Meta-analysis of the functional neuroanatomy of single-word reading: Method and validation. Neuroimage 16, 765– 780. https://doi.org/10.1006/nimg.2002.1131

Uchiyama, Y., Ebe, K., Kozato, A., Okada, T., Sadato, N., 2003. The neural substrates of driving at a safe distance: A functional MRI study. Neurosci. Lett. 352, 199–202. https://doi.org/10.1016/j.neulet.2003.08.072

Uchiyama, Y., Toyoda, H., Sakai, H., Shin, D., Ebe, K., Sadato, N., 2012. Suppression of brain activity related to a car-following task with an auditory task: An fMRI study. Transp. Res. Part F Traffic Psychol. Behav. 15, 25–37. https://doi.org/10.1016/j.trf.2011.11.002

Valentin, V. V., Dickinson, A., O’Doherty, J.P., 2007. Determining the neural substrates of goal-directed learning in the human brain. J. Neurosci. 27, 4019–4026. https://doi.org/10.1523/JNEUROSCI.0564-07.2007

van Steenbergen, H., Watson, P., Wiers, R.W., Hommel, B., de Wit, S., 2017. Dissociable corticostriatal circuits underlie goal-directed vs. cue-elicited habitual food seeking after satiation: evidence from a multimodal MRI study. Eur. J. Neurosci. 46, 1815–1827. https://doi.org/10.1111/ejn.13586

Vannest, J., Eaton, K.P., Henkel, D., Siegel, M., Tsevat, R.K., Allendorfer, J.B., Schefft, B.K., Banks, C., Szaflarski, J.P., 2012. Cortical correlates of self-generation in verbal paired associate learning. Brain Res. 1437, 104–114. https://doi.org/10.1016/j.brainres.2011.12.020

Vaquero, L., Hartmann, K., Ripollés, P., Rojo, N., Sierpowska, J., François, C., Càmara, E., van Vugt, F.T., Mohammadi, B., Samii, A., Münte, T.F., Rodríguez-Fornells, A., Altenmüller, E., 2016. Structural neuroplasticity in expert pianists depends on the age of musical training onset. Neuroimage 126, 106–119. https://doi.org/10.1016/j.neuroimage.2015.11.008

Varotto, S.F., Farah, H., Bogenberger, K., van Arem, B., Hoogendoorn, S.P., 2020. Adaptations in driver behaviour characteristics during control transitions from full-range Adaptive Cruise Control to manual driving: an on-road study. Transp. A Transp. Sci. 16, 776–806. https://doi.org/10.1080/23249935.2020.1720856

Vigneau, M., Jobard, G., Mazoyer, B., Tzourio-Mazoyer, N., 2005. Word and non-word reading: What role for the Visual Word Form Area? Neuroimage 27, 694–705. https://doi.org/10.1016/j.neuroimage.2005.04.038

Vonsattel, J.P.G., DiFiglia, M., 1998. Huntington disease. J. Neuropathol. Exp. Neurol. 57, 369–384. https://doi.org/10.1097/00005072-199805000-00001

Wan Lee, S., Shimojo, S., O’Doherty, J.P., 2014. Neural computations underlying arbitration between model-based and model-free learning. Neuron 81, 687. https://doi.org/10.1016/J.NEURON.2013.11.028

Watson, P., de Wit, S., 2018. Current limits of experimental research into habits and future directions. Curr. Opin. Behav. Sci. 20, 33–39. https://doi.org/10.1016/j.cobeha.2017.09.012

Watson, P., van Wingen, G., de Wit, S., 2018. Conflicted between goal-directed and habitual control, an fMRI investigation. eNeuro 5. https://doi.org/10.1523/ENEURO.0240-18.2018

Watson, P., Wiers, R.W., Hommel, B., De Wit, S., 2014. Working for food you don’t desire. Cues interfere with goal-directed food-seeking. Appetite 79, 139–148. https://doi.org/10.1016/j.appet.2014.04.005

Wechsler, K., Drescher, U., Janouch, C., Haeger, M., Voelcker-Rehage, C., Bock, O., 2018. Multitasking during simulated car driving: A comparison of young and older persons. Front. Psychol. 9, 910. https://doi.org/10.3389/fpsyg.2018.00910

West, R., Brown, J., 2013. Addiction, habit and instrumental learning. Theory Addict. 114–135. https://doi.org/10.1002/9781118484890.CH5

Willingham, D.B., Koroshetz, W.J., 1993. Evidence for dissociable motor skills Huntington’s disease patients. In. Psychobiology 21.

Wong, A.L., Goldsmith, J., Forrence, A.D., Haith, A.M., Krakauer, J.W., 2017. Reaction times can reflect habits rather than computations. Elife 6. https://doi.org/10.7554/eLife.28075

Wood, W., Mazar, A., Neal, D., 2021. Habits and Goals in Human Behavior: Separate but Interacting Systems. Perspect. Psychol. Sci. https://doi.org/10.31234/OSF.IO/QVRBY

Wood, W., Neal, D.T., 2007. A New Look at Habits and the Habit-Goal Interface. Psychol. Rev. 114, 843–863. https://doi.org/10.1037/0033-295X.114.4.843

Worringer, B., Langner, R., Koch, I., Eickhoff, S.B., Eickhoff, C.R., Binkofski, F.C., 2019. Common and distinct neural correlates of dual-tasking and task-switching: a meta-analytic review and a neuro-cognitive processing model of human multitasking. Brain Struct. Funct. 224, 1845–1869. https://doi.org/10.1007/s00429-019-01870-4

Wu, T., Hallett, M., Chan, P., 2015. Motor automaticity in Parkinson’s disease. Neurobiol. Dis. 82, 226–234. https://doi.org/10.1016/j.nbd.2015.06.014

Wu, T., Zhang, J., Hallett, M., Feng, T., Hou, Y., Chan, P., 2016. Neural correlates underlying Micrographia in Parkinson’s disease. Brain 139, 144–160. https://doi.org/10.1093/brain/awv319

Wunderlich, K., Dayan, P., Dolan, R.J., 2012. Mapping value based planning and extensively trained choice in the human brain. Nat. Neurosci. 15, 786–791. https://doi.org/10.1038/nn.3068

Wymbs, N.F., Bassett, D.S., Mucha, P.J., Porter, M.A., Grafton, S.T., 2012. Differential Recruitment of the Sensorimotor Putamen and Frontoparietal Cortex during Motor Chunking in Humans. Neuron 74, 936–946. https://doi.org/10.1016/j.neuron.2012.03.038

Yamada, H., Matsumoto, N., Kimura, M., 2004. Tonically Active Neurons in the Primate Caudate Nucleus and Putamen Differentially Encode Instructed Motivational Outcomes of Action. J. Neurosci. 24, 3500–3510. https://doi.org/10.1523/JNEUROSCI.0068-04.2004

Yang, Y., Zhang, J., Meng, Z.L., Qin, L., Liu, Y.F., Bi, H.Y., 2018. Neural correlates of orthographic access in Mandarin Chinese writing: An fMRI study of the word-frequency effect. Front. Behav. Neurosci. 12, 1–12. https://doi.org/10.3389/fnbeh.2018.00288

Yarkoni, T., Speer, N.K., Balota, D.A., McAvoy, M.P., Zacks, J.M., 2008. Pictures of a thousand words: Investigating the neural mechanisms of reading with extremely rapid event-related fMRI. Neuroimage 42, 973–987. https://doi.org/10.1016/j.neuroimage.2008.04.258

Zwosta, K., Ruge, H., Goschke, T., Wolfensteller, U., 2018. Habit strength is predicted by activity dynamics in goal-directed brain systems during training. Neuroimage 165, 125–137. https://doi.org/10.1016/j.neuroimage.2017.09.062

## Supplementary References

Allali, G., Montembeault, M., Brambati, S.M., Bherer, L., Blumen, H.M., Launay, C.P., Liu-Ambrose, T., Helbostad, J.L., Verghese, J., Beauchet, O., 2019. Brain structure covariance associated with gait control in aging. Journals Gerontol. - Ser. A Biol. Sci. Med. Sci. 74, 705–713. https://doi.org/10.1093/gerona/gly123

Bookheimer, S.Y., Zeffiro, T.A., Blaxton, T., Gaillard, W., Theodore, W., 1995. Regional cerebral blood flow during object naming and word reading. Hum. Brain Mapp. 3, 93–106. https://doi.org/10.1002/hbm.460030206

Leh, S.E., Ptito, A., Chakravarty, M.M., Strafella, A.P., 2007. Fronto-striatal connections in the human brain: A probabilistic diffusion tractography study. Neurosci. Lett. 419, 113–118. https://doi.org/10.1016/j.neulet.2007.04.049

Müller, V.I., Cieslik, E.C., Laird, A.R., Fox, P.T., Radua, J., Mataix-Cols, D., Tench, C.R., Yarkoni, T., Nichols, T.E., Turkeltaub, P.E., Wager, T.D., Eickhoff, S.B., 2018a. Ten simple rules for neuroimaging meta-analysis. Neurosci. Biobehav. Rev. 84, 151. https://doi.org/10.1016/J.NEUBIOREV.2017.11.012

Müller, V.I., Cieslik, E.C., Laird, A.R., Fox, P.T., Radua, J., Mataix-Cols, D., Tench, C.R., Yarkoni, T., Nichols, T.E., Turkeltaub, P.E., Wager, T.D., Eickhoff, S.B., 2018b. Ten simple rules for neuroimaging meta-analysis. Neurosci. Biobehav. Rev. https://doi.org/10.1016/j.neubiorev.2017.11.012

West, R., Brown, J., 2013. Addiction, habit and instrumental learning. Theory Addict. 114– 135. https://doi.org/10.1002/9781118484890.CH5

